# smoothclust: Spatial smoothing and boundary-density summaries for spatial transcriptomics

**DOI:** 10.64898/2026.10.26.761912

**Authors:** Lauren Rusnak, Lukas M. Weber

## Abstract

Spatial transcriptomics enables the measurement of gene expression while preserving tissue organization. A common analysis task is the identification of spatial domains, defined here as spatially coherent regions with consistent expression profiles or cell-type mixtures and represented using categorical spatial tissue maps. However, while numerous methods have been developed to identify spatial domains, interpretable summary measures to quantify spatial domain organization and tissue architecture at the sample level remain less standardized. Here, we present smoothclust, a framework for spatial domain analysis consisting of two components: a spatial-smoothing-based clustering workflow for generating spatial domain labels, and a boundary density (BD) metric that provides local and sample-level summaries of label transitions on a spatial graph. Building on established categorical spatial statistics, the raw BD metric is calculated from the proportion of neighboring spatial locations that are assigned to different domain labels, and requires only spatial coordinates and domain labels as inputs. The metric can be calculated using either our spatial-smoothing-based domain labels or labels from external clustering methods or manual annotations. Relative BD is a composition-adjusted summary, calculated by dividing raw BD by its expectation under random relabeling with the observed label counts. By quantifying label transitions, our framework translates spatial domain label maps into interpretable local and sample-level summary measures of tissue architecture, facilitating exploratory analyses and comparisons across samples and biological conditions. We demonstrate the approach through evaluations across datasets from several spatial transcriptomics platforms and simulations. The spatial-smoothing workflow and BD metric are implemented in the smoothclust R package, which is freely available from Bioconductor and GitHub.

## 1. Introduction

Spatial transcriptomics technologies measure high-dimensional gene expression together with spatial coordinates in intact tissue sections, enabling direct investigation of tissue organization in cancer, neuroscience, immunology, development, and other settings [1, 2, 3]. Technological platforms include sequencing-based and imaging-based approaches, and differ in spatial resolution, transcriptome coverage, sensitivity, and whether measurements correspond to spots, bins, segmented cells, or subcellular transcript detections [2, 4]. Computational analyses of these data consist of complex workflows integrating a number of preprocessing and downstream analyses [5, 6, 7, 8]. A key analysis task is the identification of spatial domains, defined here as spatially coherent regions with consistent expression profiles or cell-type mixtures, which are represented using categorical spatial tissue maps. Depending on the biological context, these maps may delineate anatomical regions, tumor, stromal, or immune compartments, cell-type niches, or other regions of shared molecular composition. Spatial domains may be identified algorithmically using spatially-aware clustering algorithms, or may be defined based on manual annotations, histology-derived segments, or anatomical regions. Spatial domain maps provide an interpretable intermediate representation between individual spatial locations and whole tissue sections, supporting analyses of domain composition, within-domain expression, domain-level differential expression, relationships to histology, and interfaces between tissue compartments.

Numerous computational methods have been developed for spatially-aware clustering and spatial domain identification, applying various methodological approaches. Representative approaches include Bayesian modeling of spatial neighborhoods and multi-sample clustering, spatially-aware dimension reduction and factor analysis, graph convolutional or graph attention models, spatial representation learning and integration, histology or neighborhood features, and expression smoothing followed by clustering [9, 10, 11, 12, 13, 14, 15, 16, 17, 18]. These methods differ in the modalities of information they use, the scale and strength of spatial regularization, the type and granularity of labels they produce, and whether they are designed for individual sections or multiple samples. Benchmarking studies have emphasized that performance depends on tissue structure, technological platform, spatial resolution, available annotations, sample-integration strategy, and the downstream scientific question, with no single method performing uniformly best across settings [19, 20].

Multi-sample spatial transcriptomics studies introduce an additional layer of challenges for analysis by creating a need to make results comparable across tissue sections, individuals, or biological conditions [20]. One strategy is to analyze multiple sections jointly by integrating expression spaces, aligning tissue sections, or fitting models that directly accept multiple sections as input. This includes general single-cell integration approaches such as Harmony [21] and others, as well as spatially-aware methods for alignment, integration, and multi-sample domain detection [22, 10, 12, 15, 23, 18]. These approaches can borrow information across samples, reduce the need for post hoc matching of section-specific labels, and support comparisons in a common embedding or label space. However, they also require choices or assumptions about batch correction, correspondence of tissue structures and labels, spatial alignment, comparable resolution, and whether sample-specific biological variation should be preserved or removed.

A complementary strategy is to calculate sample-level summary measures after categorical labels have been obtained, whether from independent single-section analyses, an integrated multi-sample workflow, histology-derived segmentation, published computational annotations, or manual annotations. When informative summary measures are available, this approach can provide a simpler and interpretable route for exploratory visualization, comparisons across biological groups or conditions, or association analyses with clinical covariates. While spatial domain detection has received substantial methodological attention, the development and interpretation of summary measures for quantifying tissue architecture at the sample level remain less standardized. Most domain detection workflows return a spatial map of labels, which is visually informative but difficult to compare systematically across large cohorts or experimental conditions. Summary-based analyses can complement joint clustering and integration without necessarily requiring alignment of individual spatial locations across sections.

Existing spatial analysis frameworks provide several approaches to this problem. These include multi-view sample representations in scFeatures [24], cell-type colocalization, neighborhood enrichment, and spatial interaction analyses in frameworks such as Squidpy, SPIAT, and spicyR [6, 25, 26], multiscale diversity summaries in MESA [27], and tissue representations based on persistent local relationship patterns in Kasumi [28]. HTA provides an entropy-based sample-level heterogeneity summary by averaging local heterogeneity across spatial regions [29]. CellCharter additionally characterizes spatial cluster components using shape-related measures [30]. These approaches describe different aspects of tissue architecture, and the choice of summary depends on the scientific question; no single spatial summary fully establishes clustering accuracy or biological validity [31, 32, 33, 34].

Here, we develop a spatial domain boundary density (BD) metric as a simple label-source-agnostic measure of label-transition frequency within a tissue section. Given spatial coordinates and a domain label for each spatial location, the raw BD metric quantifies the proportion of local nearest-neighbor relationships that connect locations assigned to different labels. Intuitively, at each spatial location, this represents the proportion of neighboring spatial locations that are assigned to different domain labels. This quantity is a normalized discordant join count and connects directly to classical categorical spatial statistics and graph homophily [35, 36, 37]. Spatial autocorrelation, pair-specific joins, and local indicators of spatial association provide related descriptions of spatial structure [38, 39, 40], with relevant implementations including stJoincount [41] and Voyager [42].

Related recent approaches include SHouT [43], inter-label connectivity in Spatialproteomics [44], a spatial-connectivity criterion used for benchmarking by [45], neighborhood entropy analyses provided by ClustSIGNAL [18], and stEDGE [46]. SHouT reports global edge homophily, defined as the fraction of spatial-graph edges connecting cells of the same type, and global normalized entropy of the sample-wide label composition as sample-level scores [43]. On an identical spatial graph, sample-level raw BD is the complement of global homophily, although the default graph definitions and intended label maps differ. At the local level, SHouT’s egophily measures agreement between a focal cell and cells in its neighborhood, making it closely related to the complement of local BD. Spatialproteomics [44] defines inter-label connectivity as the fraction of a cell’s neighbors assigned to a different cell-type label, and its average across cells for sample comparisons; with matching labels and spatial neighborhoods excluding the focal cell, the local measure and equally-weighted average are equivalent to local and sample-level raw BD. [45] use a spatial-connectivity criterion for benchmarking niche identification that averages the fraction of neighboring cells assigned to the same domain label as the focal cell; under matching labels, spatial neighborhoods, and averaging conventions, this is equivalent to one minus sample-level raw BD. ClustSIGNAL calculates cell-specific Shannon entropy from the initial-subcluster composition of each neighborhood and uses these values to control adaptive expression smoothing; the resulting neighborhood entropy distributions can also be summarized at the sample level [18]. These entropy summaries quantify neighborhood diversity, whereas local BD quantifies disagreement between a focal location’s label and the labels of its neighbors. stEDGE uses a consensus-based boundary signal to guide spatial domain reconstruction and hierarchical organization [46]. With matching neighborhoods and exclusion of self-relations, its initial boundary signal can be expressed as local raw BD averaged over an ensemble of labelings. However, stEDGE uses this signal to reconstruct domains, whereas we use BD to summarize existing label maps.

An optional composition adjustment to the raw BD metric produces relative BD, calculated by dividing raw BD by its expectation under random relabeling with the observed label counts. This expectation can be calculated analytically from the label counts. Relative BD expresses observed transition frequency relative to a section-specific reference, complementing the direct summary provided by raw BD. It also has a connection to Geary’s *C* [39], connecting BD to established spatial statistics: for binary labels and matching spatial weights, relative BD equals Geary’s *C*; for multiple labels, it is a variance-weighted average of Geary’s *C* calculated for the corresponding label-indicator variables. The adjustment accounts for label composition through the random-label expectation but does not make relative BD independent of composition, label definitions, or spatial scale. Comparisons therefore require comparable labels, spatial graphs, and sampling units.

We present smoothclust, a framework for spatial domain analysis consisting of two components: a spatial-smoothing-based clustering workflow for generating spatial domain labels, and a BD metric that provides local and sample-level summaries of label transitions on a spatial graph. The BD metric requires only spatial coordinates and categorical labels as inputs, and can be applied to labels from smoothclust, external clustering methods, published annotations, or manual annotations. Low raw BD indicates relatively few local label transitions, while high raw BD indicates frequent local transitions, which may reflect dense interfaces between tissue compartments, intermixed biological structures, fragmented domains, or noisy label assignments. Our contribution combines an accessible R/Bioconductor implementation with analyses of datasets from several spatial transcriptomics technologies demonstrating how BD summaries depend on label definitions, spatial scale, and the datasets being studied. By quantifying spatial domain label transitions, our framework translates categorical spatial maps into interpretable local and sample-level summary measures of tissue architecture, facilitating exploratory analyses and sample-level comparisons.

## 2 Results

### 2.1 BD metric summarizes GBM/glioma tissue organization

We first applied smoothclust independently to 19 Greenwald et al. GBM/glioma Visium sections [47] (Supplementary Table S1) and compared the resulting labels with a matched non-spatial *k*-means baseline (Figure 1). Both workflows used the same retained spots and input genes, normalization and transformation procedures, and downstream analysis settings. Variable genes and principal components were estimated separately from smoothed and unsmoothed expression. This comparison therefore evaluated the effect of including smoothing in the complete clustering workflow.

**Figure 1:**
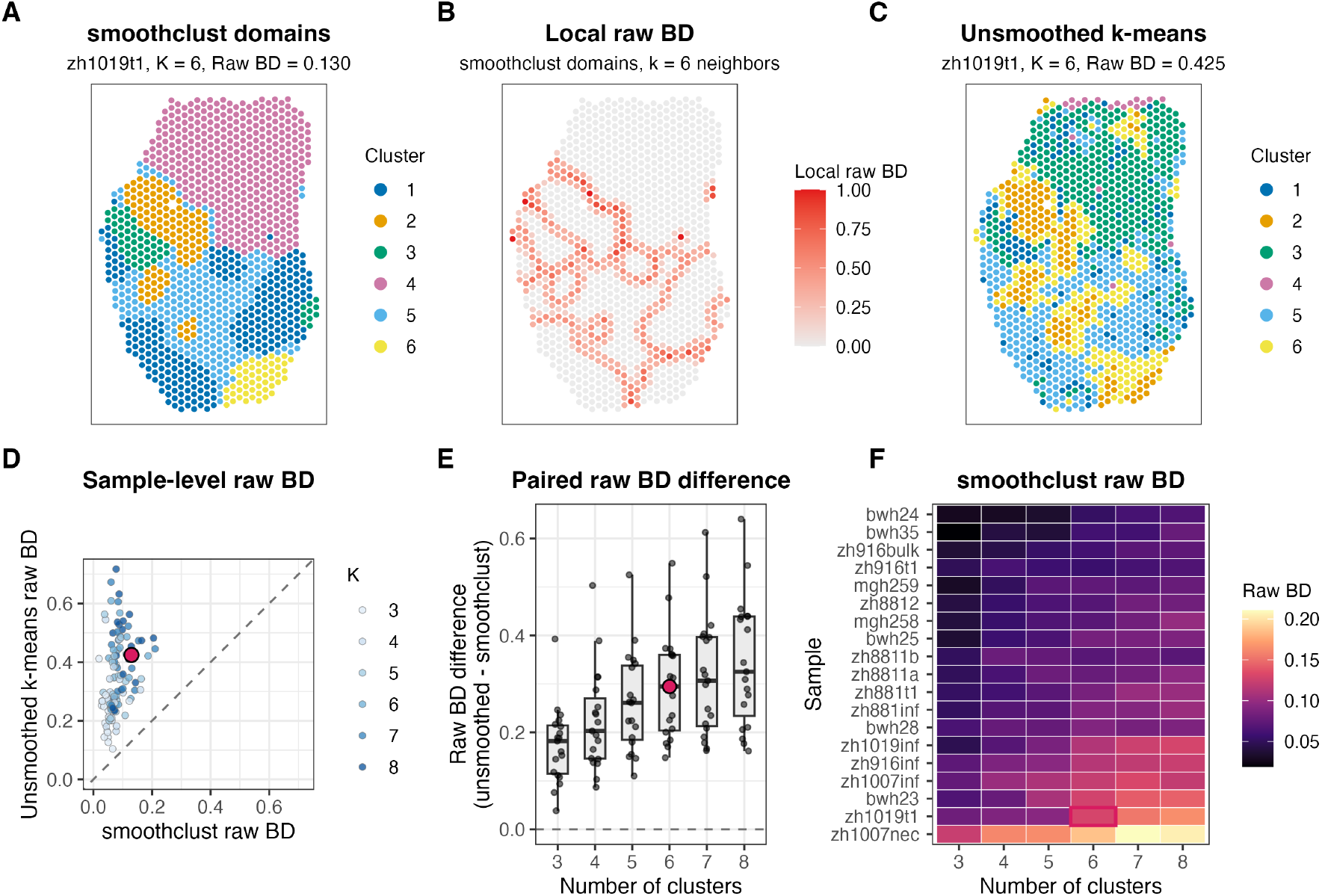
The BD metric summarizes the effect of spatial smoothing on GBM/glioma label maps. (A) smoothclust domain labels for section zh1019t1 with *K* = 6. (B) Corresponding local raw BD map for the smoothclust labels. (C) Domain labels generated using the matched non-spatial *k*-means baseline without smoothing. (D) Paired sample-level raw BD values across 19 sections and *K* = 3, … , 8; the dashed line indicates equality. (E) Paired differences in raw BD (baseline minus smoothclust), grouped by cluster number. (F) Sample-level raw BD for smoothclust across sections and cluster numbers. The highlighted points and tile in panels D–F mark zh1019t1 at *K* = 6. All BD calculations used *k* = 6. Feature selection and PCA were performed separately for the smoothed and unsmoothed workflows.

Local raw BD highlighted interfaces between domain labels for smoothclust (Figure 1A–B), while the matched non-spatial baseline showed more frequent label transitions (Figure 1C). In section zh1019t1 at *K* = 6, sample-level raw BD was 0.130 for smoothclust and 0.425 for the baseline. Across all 19 sections and *K* = 3, … , 8, the baseline had higher raw BD in all 114 matched comparisons (Figure 1D– E). At *K* = 6, mean raw BD was 0.091 for smoothclust and 0.386 for the baseline. Sample-level raw BD values for smoothclust across sections and cluster numbers are summarized in Figure 1F. These results demonstrate the reduction in label transitions after spatial smoothing.

Lower raw BD values after smoothing were also observed with kernel and *k*-nearest-neighbor smoothing (Supplementary Figure S1). Increasing the smoothing scale generally lowered raw BD (Supplementary Figure S2). In contrast, holding labels fixed and increasing the BD neighborhood size broadened local interface signals and generally raised the sample-level value (Supplementary Figure S3). These analyses illustrate the distinct effects of smoothing scale and the neighborhood used to summarize the resulting labels.

### 2.2 Simulations illustrate the composition adjustment and over-smoothing

Composition simulations on a 30 × 30 grid illustrated the distinction between raw and relative BD (Supplementary Figure S4). Random arrangements at 10:90, 30:70, and 50:50 label compositions had different raw BD expectations, while relative BD was centered near 1, with means ranging from 1.000 to 1.002 across the three compositions. With counts fixed at 450:450, relative BD increased from 0.034 for two coherent halves to 0.391 and 0.836 after exchanging labels at 10% and 30% of locations, respectively, 0.976 for a random arrangement, and 1.931 for a checkerboard. At this fixed composition, raw and relative BD differ only by a constant scaling.

Straight-interface maps with 10:90, 30:70, and 50:50 compositions further illustrated the role of the random-label expectation. With the graph, interface orientation, and interface extent held fixed, raw BD was approximately 0.017 in all three maps, while relative BD was 0.096, 0.040, and 0.034, respectively. Thus, similar observed transition frequencies can yield different relative BD values because the expected frequencies differ. The adjustment therefore does not make relative BD independent of composition in spatially structured maps.

An additional over-smoothing simulation illustrated why lower BD does not necessarily indicate better domain recovery (Supplementary Figure S5). For two broad domains and a small island, recovery was highest at a uniform-smoothing bandwidth of 4% of the coordinate range (ARI 0.973; island F1 0.872). At 30%, ARI fell to 0.663 and island F1 to 0.105 even though raw and relative BD were lower.

This simulation illustrates how excessive smoothing can erase a small domain while further reducing label transitions; the best-performing bandwidth is specific to this example.

### 2.3 BD metric summarizes external spatial clustering labels or manual annotations

We next applied BD to domain labels that we generated using three external spatial clustering algorithms. Across the 19 sections for the GBM/glioma Visium dataset, median sample-level raw BD was 0.103 for spatial clustering labels generated using BayesSpace [9], 0.085 for BANKSY [17], and 0.155 for ClustSIGNAL [18], compared with 0.071 for smoothclust clustering and 0.343 for the non-spatial baseline (Supplementary Figure S6). BayesSpace, BANKSY, smoothclust, and the baseline used *K* = 5, while ClustSIGNAL used its default labeling settings and returned a variable number of labels [18]. For section zh1019t1, the domain labels generated by BayesSpace, BANKSY, and ClustSIGNAL are shown in Figure 2A–C, with the corresponding local raw BD maps highlighting label transitions in Figure 2D–F. These analyses illustrate the flexibility of BD as a summary measure that can be calculated from any source of categorical spatial domain labels.

**Figure 2:**
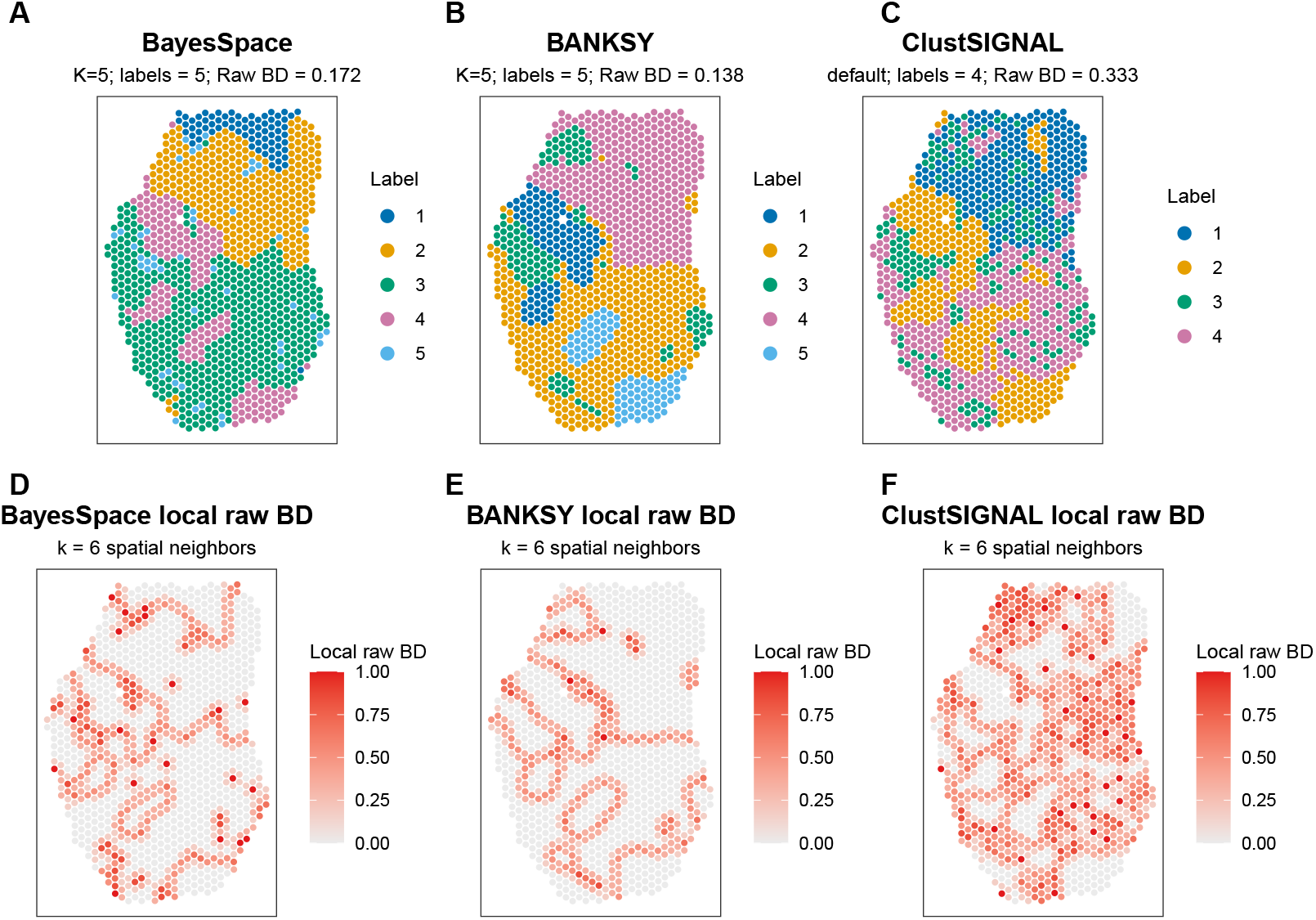
The BD metric summarizes labels generated using external spatial clustering algorithms. (A–C) Domain labels obtained by applying BayesSpace [9], BANKSY [17], and ClustSIGNAL [18], respectively, to section zh1019t1. (D–F) Corresponding local raw BD maps on a common 0–1 scale. Colors and cluster numbers do not imply matched biological identities across methods. BayesSpace and BANKSY used *K* = 5; ClustSIGNAL used its default settings and returned four labels in this section. All BD calculations used *k* = 6.

We also applied BD directly to four sets of published annotation labels from Greenwald et al. [47], illustrating the use of BD with existing label maps rather than labels generated by independent clustering. Across the 13 sections with these annotations, coarse organization-zone labels had lower transition frequencies than dominant-metaprogram labels and deposited layer annotations (Supplementary Figure S7). Median relative BD was 0.117 for coarse organization zones, 0.162 for malignancy-refined zones, 0.771 for dominant metaprograms, and 0.746 for layer labels. Metaprogram and layer relative BD were negatively associated with the published structured-zone fraction (Spearman *ρ* = −0.90 and −0.86, respectively). Since the organization zones were derived from local coherence of the metaprogram assignments, these associations show agreement between related descriptions of tissue organization.

We also examined regional variation within individual tumors by comparing metaprogram composition and relative BD across sections from the same tumor. Across 12 regional sections from four tumors, we observed substantial within-tumor differences, particularly for ZH881 and ZH916 (Supplementary Figure S8).

### 2.4 BD metric provides a sample-level summary of myocardial organization

We next applied BD to a different biological setting to illustrate its use for descriptive comparisons of tissue organization across sample categories. We calculated BD using the nine published cell-type-niche labels available from the myocardial-infarction Visium dataset of Kuppe et al. [48], across 28 tissue sections from 20 patients. Median relative BD was 0.923 in myogenic-enriched, 0.656 in ischaemic-enriched, and 0.737 in fibrotic-enriched sections (Figure 3A). All relative BD values were below 1, indicating fewer label transitions than expected under random relabeling with the observed counts in each section, as illustrated by the comparison of observed and expected raw BD (Figure 3B). Local raw BD maps showed the transitions contributing to representative sample-level values (Supplementary Figure S9). Higher relative BD in myogenic-enriched sections indicates more frequent transitions among the published cell-type-niche labels relative to their section-specific composition references. This pattern may reflect interspersion of compositionally distinct niches within broadly myogenic tissue, whereas some injured sections may contain larger patches dominated by particular niche labels.

**Figure 3:**
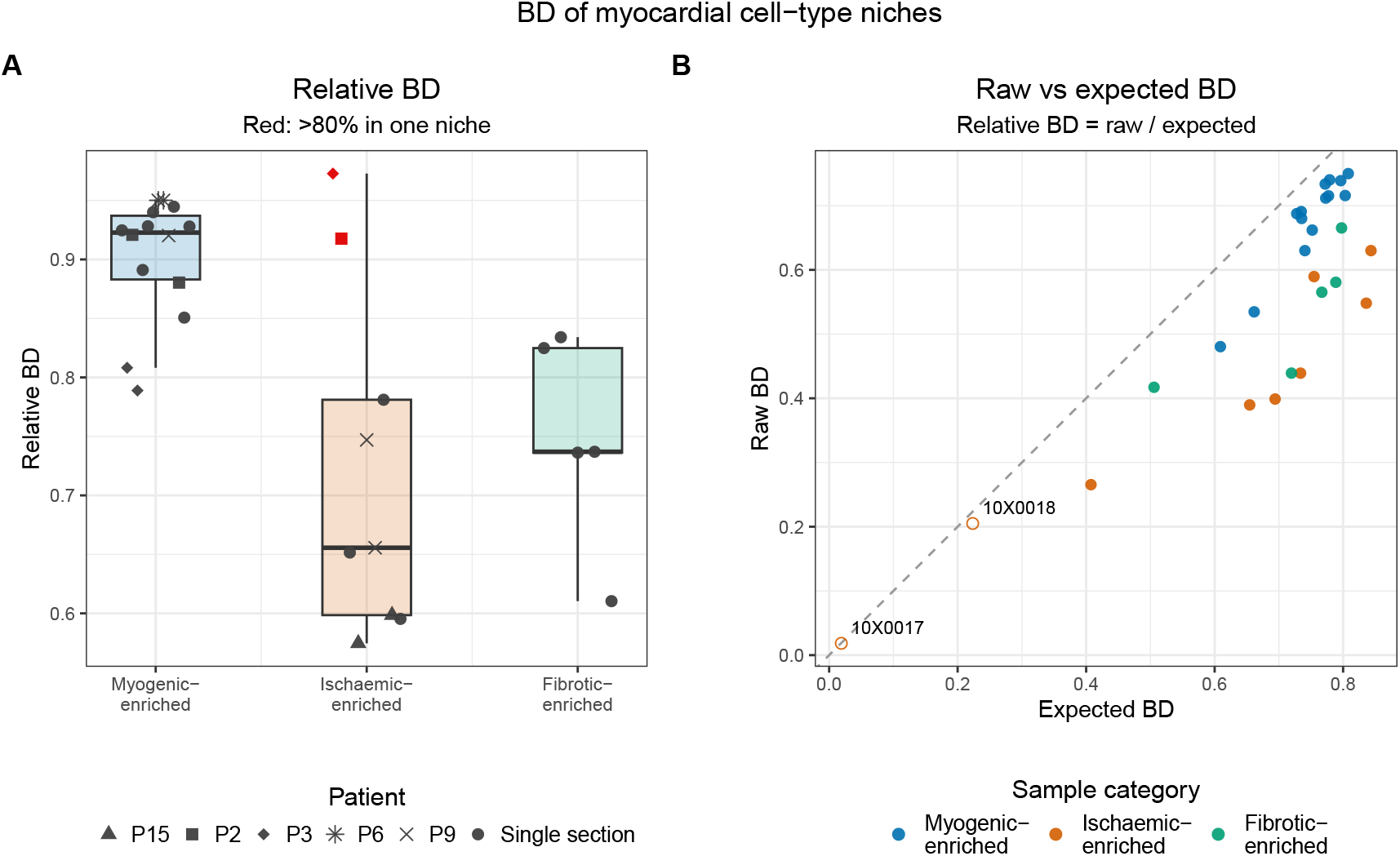
Relative BD summarizes published myocardial cell-type-niche organization. (A) Relative BD for the nine published cell-type-niche labels from Kuppe et al. [48] across 28 sections from 20 patients, grouped by the author-defined tissue categories: myogenic-enriched (14 sections), ischaemic-enriched (nine), and fibrotic-enriched (five). Each point is one section; boxes show medians and interquartile ranges and are colored by sample category. Shapes identify patients with multiple sections; circles denote patients with a single section. Red points mark two ischaemic sections in which one niche occupied more than 80% of spots. (B) Observed sample-level raw BD plotted against expected raw BD under random relabeling with the observed label counts in each section; the dashed line denotes equality. Point colors match the box colors in (A), and open circles mark the same two flagged sections. All BD calculations used *k* = 6. Group summaries are descriptive and section-weighted.

Two ischaemic sections were dominated by a single niche, occupying 99.0% and 87.8% of spots (red points in Figure 3A and open circles in Figure 3B; composition profiles in Supplementary Figure S10). Excluding these sections changed the ischaemic median relative BD only slightly, from 0.656 to 0.652. This illustrates how BD can provide section-level summaries even for sections with strongly imbalanced label compositions. Comparisons across the author-defined tissue categories were descriptive and section-weighted, with repeated patient identities indicated in Figure 3A.

To assess sensitivity to neighborhood size, we recalculated relative BD using *k* = 18 and *k* = 36 while holding the published niche labels fixed. Section-level rankings remained strongly correlated with those at *k* = 6 (Spearman *ρ* = 0.989 and 0.984, respectively; Supplementary Figure S11), with median absolute changes of 0.025 and 0.039. Category medians retained the ordering myogenic-enriched *>* fibrotic-enriched *>* ischaemic-enriched.

### 2.5 BD metric provides annotation-specific cortical summaries

To further investigate the effects of annotation granularity on BD, we analyzed a human frontal-cortex Visium dataset from Miyoshi et al. [49]. The dataset includes published annotations generated with BayesSpace [9] that define nine cortical and white-matter regions across 39 tissue sections. Using these labels, we obtained relative BD values ranging from 0.127 to 0.312, with a median of 0.198 (Supplementary Figure S12). Plotting section-level values alongside label proportions provides a joint view of spatial organization and composition. Complete annotation and local raw BD maps are shown in Supplementary Figure S13.

For the original nine-region annotation, we also assessed sensitivity to BD neighborhood size. Relative BD rankings at *k* = 18 and *k* = 36 remained strongly correlated with those at *k* = 6 (Spearman *ρ* = 0.965 and 0.925, respectively), although values shifted substantially (median absolute changes of 0.078 and 0.145, respectively; Supplementary Figure S11).

To investigate the effects of annotation granularity, we then collapsed WM1–WM3 into a single white-matter label. The original and merged annotations, together with their corresponding local raw BD maps for a representative section, are shown in Figure 4A–B. Across all 39 sections, raw BD decreased in 38 and was unchanged in one, with a median paired change (merged minus original) of −0.0254 (Figure 4C). Merging labels also decreased or left unchanged the random-label expectation (Figure 4D), since pairs assigned to different white-matter labels become concordant when at least two of the merged labels are present. Median relative BD decreased to 0.175, and section rankings changed (Spearman *ρ* = 0.591; Figure 4E). Relative BD decreased in 37 sections, was unchanged in one, and increased in one, with a median paired change of −0.019. Since relative BD is the ratio of observed to expected BD, it can increase when the expectation decreases proportionally more than the observed value. These results illustrate how label granularity affects both the magnitude and ordering of sample-level summaries.

**Figure 4:**
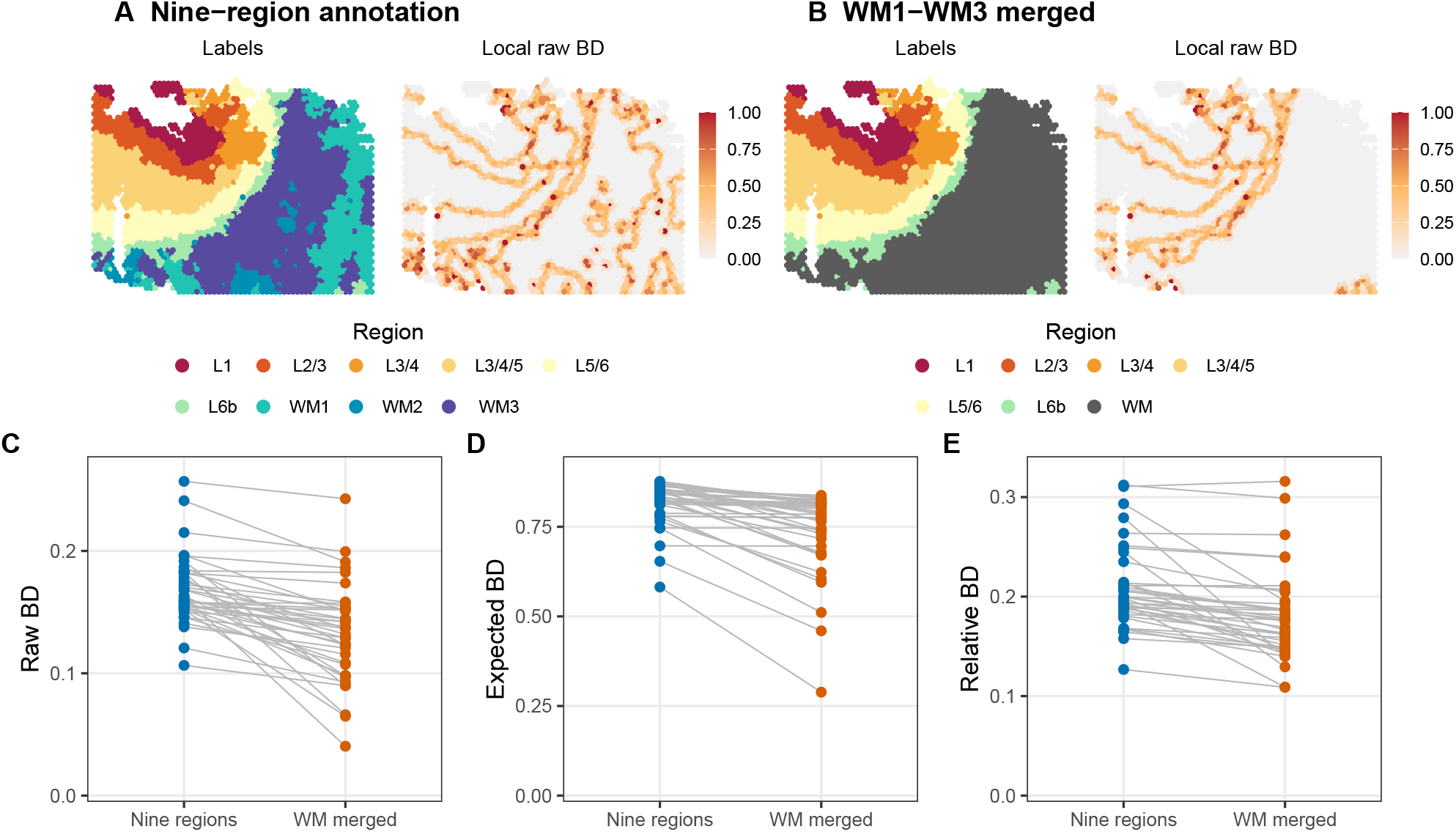
Raw, expected, and relative BD summarize the effects of cortical annotation granularity. (A) Published nine-region BayesSpace annotation and corresponding local raw BD map for a representative Miyoshi frontal-cortex Visium section (`Nov_24_2021_VisiumHuman_12`), selected as the 20th of 39 sections ordered by increasing relative BD under the original annotation. (B) The same section after merging WM1–WM3 into a single white-matter label, with the corresponding local raw BD map. (C) Paired sample-level raw BD values across all 39 sections under the original and merged annotation definitions. (D) Corresponding expected raw BD under random relabeling with the observed label counts, calculated analytically for each section and label definition. (E) Corresponding relative BD, calculated as raw BD divided by expected raw BD. Lines in C–E connect values from the same section. Retained spots, spot order, transformed coordinates, and directed *k* = 6 neighbor-search settings were the same for both definitions; only the label definition changed. Local raw BD maps in A–B use a common 0–1 color scale.

To examine BD summaries using manual anatomical annotations, we also analyzed a human dorsolateral prefrontal cortex (DLPFC) Visium dataset from Maynard et al. [50]. Using the published cortical-layer and white-matter annotations for 12 sections from three donors, relative BD ranged from 0.0666 to 0.1180. The median absolute difference between six pairs of directly adjacent sections was 0.00125, with a maximum of 0.00601 (Supplementary Figure S14). These observations provide a descriptive assessment of agreement in section-level relative BD between adjacent sections. Complete manual annotation and corresponding local raw BD maps are shown in Supplementary Figure S14.

### 2.6 High-resolution workflows and computational runtime

To assess computational feasibility and runtime on large, high-resolution spatial transcriptomics datasets, we analyzed Visium HD and Xenium data from human colorectal cancer tissue sections generated by de Oliveira et al. [51]. We applied the smoothing, clustering, and BD workflow to process 545,850 nonempty 8 *µ*m Visium HD bins and 339,681 nonempty Xenium cells from one P2CRC section per platform. The Visium HD workflow used 1,000 selected variable genes, while the Xenium workflow used all 422 imported genes. For both platforms, PCA and *K* = 15 *k*-means clustering were fitted on separate 30,000-location subsets, followed by full-data projection and assignment (see Supplementary Methods). Relative BD for the smoothclust-derived labels was 0.181 for Visium HD and 0.186 for Xenium; for the 15 distributed Visium HD labels, it was 0.364 (Supplementary Figure S15). These examples demonstrate the feasibility of applying the workflow to full high-resolution sections using subset-based fitting for PCA and clustering. The BD values describe each label map under its platform-specific graph and sampling units.

Observed runtimes for the core workflows were 54.0 seconds for Visium HD and 16.2 seconds for Xenium (Supplementary Table S2; Supplementary Figure S16). For the 19-section GBM analysis at *K* = 5, the corresponding runtime was 33.9 seconds, reconstructed from component timers. We also compared runtimes across clustering methods in the GBM dataset. Under the evaluated settings, smoothclust, BANKSY, ClustSIGNAL, and BayesSpace required 33.9 seconds, 31.1 seconds, 52.2 seconds, and 1.72 hours, respectively (Supplementary Figure S16). Further timing details are provided in Supplementary Table S2 and Supplementary Figure S16.

## 3. Discussion

The smoothclust workflow and package provide a framework for translating categorical spatial maps into local and sample-level summaries of tissue organization. The framework combines a spatial-smoothing-based clustering approach for generating domain labels with a BD metric that can be calculated independently using labels from other algorithms, published annotations, or manual annotations. A key strength of BD is that it provides a simple, interpretable sample-level summary measure for exploratory comparisons across tissue sections, while retaining a direct connection to local label transitions and offering an optional composition adjustment based on expected transitions under random relabeling with the observed label counts. Our contribution combines an accessible R/Bioconductor implementation with analyses across spatial transcriptomics technologies, illustrating how BD summaries depend on label composition, label definitions, and spatial scale.

The applications illustrate several distinct uses of the BD metric. The GBM/glioma analyses quantify the effect of spatial smoothing on label organization and demonstrate the use of BD with labels generated by smoothclust and external clustering algorithms, as well as published tissue annotations, including comparisons among regional sections from the same tumor. The myocardial example uses published cell-type-niche labels for exploratory comparisons across author-defined tissue categories, including sections with strongly imbalanced label compositions. The cortical examples show how merging related annotation labels can change both the magnitude and ranking of section-level BD summaries, while manual anatomical annotations provide a separate example of agreement in relative BD between adjacent sections. Together, these analyses illustrate how BD can support different descriptive questions across datasets and biological contexts.

Raw and relative BD provide complementary summaries of tissue organization and should be interpreted together. Raw BD describes the observed frequency of label transitions, whereas relative BD expresses this frequency relative to its expectation under random relabeling with the observed label counts. Both quantities are informative: differences in label composition may themselves be biologically meaningful, and the adjustment does not make relative BD independent of composition in spatially structured maps. Both measures also depend on label definitions, graph construction, and spatial scale, so comparisons across tissue sections require comparable labels, neighborhoods, and sampling units. BD therefore provides an interpretable summary of overall label transitions at the chosen spatial scale, complementing more detailed analyses of boundary geometry and interactions between specific labels.

Spatial smoothing introduces a separate choice of scale. The strength of spatial smoothing should be selected with the biological context and intended spatial resolution in mind, since excessive smoothing may obscure small but biologically meaningful structures, as illustrated by the over-smoothing simulation. The default uniform-radius smoother uses a bandwidth proportional to each section’s coordinate extent, so the same nominal bandwidth can correspond to different physical smoothing radii across sections. Where a common physical scale is important, section-specific bandwidths can be chosen accordingly; an absolute-distance bandwidth option could further simplify this choice. Tissue-constrained smoothing neighborhoods may also help avoid averaging expression across gaps within tissue sections. An important future direction is to apply BD in larger spatial transcriptomics studies with biological replication, enabling investigations of associations between tissue organization and clinical or experimental outcomes. Such analyses could use statistical models that account for patient or donor identity, distinguishing variation between individuals from variation among sections sampled from the same individual. BD could also serve as an exploratory sample-level diagnostic to identify sections with unusual spatial organization that warrant further investigation. Further evaluation of sensitivity to annotation uncertainty, graph construction, and tissue sampling would help establish the reliability of BD summaries in these settings.

Overall, smoothclust provides a transparent and reusable framework for characterizing spatial domain organization and tissue architecture through interpretable local and sample-level summaries of label transitions. The Visium HD and Xenium analyses further demonstrate the feasibility of applying the workflow to tissue sections containing hundreds of thousands of spatial locations on a laptop. By linking local visualization with sample-level summaries and illustrating how label composition, label definitions, and spatial scale influence BD, the framework supports exploratory analyses and comparisons of tissue organization across sections.

## 4 Methods

### 4.1 Framework and spatial smoothing

The smoothclust framework consists of two components: a spatial-smoothing-based clustering workflow and the boundary density (BD) metric. The clustering workflow normalizes gene expression counts, smooths normalized expression profiles across spatial neighbors, and log-transforms them before down-stream clustering, while BD summarizes label transitions on a spatial graph. BD provides local and sample-level summaries for each tissue section and can be calculated independently of the smoothing workflow, using labels from smoothclust, external clustering algorithms, published annotations, or manual annotations.

Let 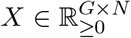 contain expression counts for *G* genes and *N* spatial locations, and let *S* ∈ ℝ^*N ×*2^ contain the matching spatial coordinates. Location- and gene-level filtering are performed before calling `smoothclust()`. If *q*_*j*_ *>* 0 is the library total used for normalization at location *j*, the size factor, rescaled to have mean one, and normalized expression are

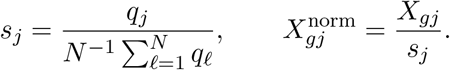

Normalization precedes smoothing so that neighboring locations contribute according to spatial weights rather than their library totals.

Spatial smoothing uses a sparse column-normalized weight matrix *W* ∈ ℝ^*N ×N*^ , with *w*_*ij*_ ≥ 0 and ∑_*i*_ *w*_*ij*_ = 1, to calculate

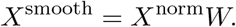

Here, *w*_*ij*_ is the contribution of location *i* to the smoothed profile at location *j*. Averaging is performed on the linear normalized-expression scale. For the default uniform-radius smoother, let *R* be the larger horizontal or vertical range of the supplied coordinates and *h* = *bR*. Each profile is replaced by the equally weighted average of itself and all other locations within Euclidean distance *h*; the default bandwidth is *b* = 0.05. The alternative kernel smoother uses exponentially decaying weights with length scale *h*, omitting unnormalized weights below *τ* = 0.05. The *k*-nearest-neighbor smoother equally weights the focal location and *k*_*s*_ neighbors, with default *k*_*s*_ = 18. The smoothing-neighbor count *k*_*s*_ is distinct from the BD neighborhood size *k*. Full weight definitions are provided in Supplementary Methods.

The default post-smoothing transformation is

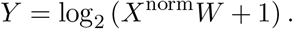

Feature selection, dimension reduction, and clustering follow the dataset-specific workflows described below, separating smoothing from the choice of clustering algorithm. Where the smoothing workflow was applied, normalization, smoothing, and clustering were performed separately within each tissue section. Analyses of published or manual annotations calculated BD directly from the supplied labels.

### 4.2 Boundary density metric

The BD metric is a graph-based summary measure of label transitions in a specified categorical label map. Let *c*_*i*_ be the label at location *i*, and let *N*_*i*_ be its spatial neighborhood, excluding *i*, with *d*_*i*_ = |*N*_*i*_| *>* 0. The local raw BD value is

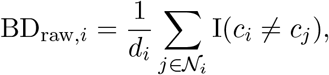

and the sample-level raw BD value is

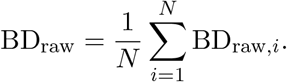

The `boundary_density()`function uses a directed Euclidean *k*-nearest-neighbor graph, with *d*_*i*_ = *k* for every location and default *k* = 6. Raw BD ranges from 0, when all neighboring labels agree, to 1, when every directed relation joins different labels.

Nearest neighbors were identified using `BiocNeighbors::findKNN()`with default search settings, independently of the domain labels. When equidistant candidates spanned the *k*-th-neighbor cutoff, the subset returned by the neighbor-search implementation was retained. Ties that only change the ordering of retained neighbors do not affect BD.

Raw BD measures label-transition frequency on a spatial graph rather than physical boundary length. Local BD highlights locations with frequent label transitions, while the sample-level mean weights each spot, bin, or cell equally. The default *k* = 6 neighborhood approximates immediate neighbors in the interior of a complete Visium lattice, although nearest-neighbor graphs may span tissue gaps or sparsely sampled regions. Comparisons therefore require comparable label definitions, graph construction, spatial scale, and sampling units.

Because raw BD also depends on label composition, we provide an optional adjustment that produces relative BD. For *L* observed labels, let *n*_ℓ_ be the number of locations carrying label ℓ, out of *N* total locations. The expected raw BD under random relabeling with the observed label counts is

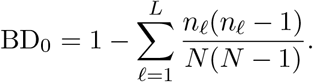

The spatial locations, graph, and label counts are held fixed while labels are reassigned uniformly among locations. Under this scheme, every ordered pair of distinct locations has the same probability of carrying different labels, yielding the expression above without requiring permutation simulations. For *L* equally abundant labels, the expectation is *N* (1 − 1*/L*)*/*(*N* − 1), and therefore increases with the number of labels at fixed *N* . More generally, the expectation depends on the complete label composition rather than the number of labels alone. Splitting a label into nonempty categories increases the expectation, whereas merging labels decreases it whenever at least two of the merged labels are present.

Relative BD is

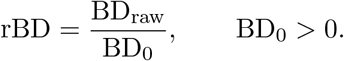

Values below, near, or above 1 indicate fewer, similar, or more label transitions than expected under random relabeling. Relative BD is undefined for single-label samples and may be unstable when expected raw BD is close to zero; a value near 1 does not establish spatial randomness.

Raw and relative BD provide complementary summaries: raw BD describes observed transition frequency, while relative BD expresses it relative to the random-label expectation. The adjustment accounts for label composition through this expectation but does not make relative BD independent of composition, label definitions, or spatial scale. Raw BD, expected raw BD, and relative BD should therefore be considered together.

### 4.3 Computational implementation and runtime

Analyses used R version 4.6.1 and smoothclust version 1.9.6. The `smoothclust()`function accepts a `SpatialExperiment`object [52] or an expression matrix with matching coordinates, uses sparse matrix operations and `BiocNeighbors`parallelization, and returns processed expression values with unchanged dimensions and names. For `SpatialExperiment`inputs, the processed values are added as a new assay and preprocessing settings are recorded in object metadata.

The `boundary density()`function operates independently on spatial coordinates and categorical labels. Labels must be nonmissing, and the neighbor count must satisfy 1 ≤ *k < N* . Its default output contains local and sample-level raw BD values; `adjust = TRUE`additionally returns expected raw BD and relative BD values at the sample level. Input handling and normalization are summarized in Supplementary Methods; detailed input requirements are provided in the R package documentation.

Runtime was measured on a 2023 MacBook Pro with an Apple M3 Max processor and 36 GB memory. Timing scope, requested threads, and the distinction between component-derived and directly observed timings are described in Supplementary Table S2 and Supplementary Figure S16.

### 4.4 Datasets and empirical analysis design

We analyzed publicly available Visium, Visium HD, and Xenium datasets using labels generated by clustering methods or obtained from published computational or manual annotations (Supplementary Table S1). BD was calculated separately for each tissue section. Repeated sections or regions from the same donor were identified in descriptive comparisons and were not treated as independent biological replicates. We used BD to characterize tissue organization in datasets from several platforms, interpreting values within each analysis in the context of differences in graph scale, spatial resolution, sampling units, and label granularity. Detailed preprocessing, feature selection, annotation matching, and coordinate transformations are provided in Supplementary Methods.

#### 4.4.1 Greenwald GBM/glioma Visium analyses

Greenwald et al. GBM/glioma Visium data were imported from GEO accession GSE237183 [47]. The clustering cohort comprised 13 GBM and six IDH-mutant glioma sections. After spot- and gene-level filtering, 34,699 of 39,566 spots and a shared set of 14,301 genes were retained across 19 sections.

Each section was analyzed independently. The primary workflow used uniform-radius smoothing with *b* = 0.05, followed by variable-gene selection, PCA with 50 components, and *k*-means clustering with 25 random starts for *K* = 3, … , 8. A matched baseline omitted smoothing before log transformation. Both workflows used the same normalization and downstream analysis procedures, with variable genes and principal components estimated separately from the smoothed and unsmoothed expression matrices.

Local and sample-level BD were calculated using *k* = 6. Sensitivity analyses compared uniform, kernel, and nearest-neighbor smoothing; varied smoothing parameters; and evaluated BD neighborhood sizes from *k* = 2 to 36 while holding labels fixed. Agreement between clusterings obtained using different smoothing methods was assessed using the adjusted Rand index (ARI).

For the external-method comparison, BayesSpace and BANKSY were run with five clusters per section, while ClustSIGNAL used its default labeling settings and returned a variable number of clusters [9, 17, 18]. All methods used the same retained spots and shared gene set. Raw BD was calculated for each resulting label map using *k* = 6.

We also analyzed four sets of published Greenwald annotation labels available for 13 GBM sections from five tumors: coarse organization zones (`org1`), malignancy-refined organization zones (`org2`), dominant metaprograms (`mp`), and deposited layer annotations (`layer`). Author-provided labels were retained as deposited. BD was calculated using *k* = 6 on spots with nonmissing labels for each field, with relative BD used for sample-level comparisons across annotation types.

Finally, we compared metaprogram composition and relative BD for metaprogram and layer annotations descriptively across 12 regional sections from four tumors, including five sections from tumor ZH881.

#### 4.4.2 Kuppe myocardial-infarction Visium analyses

Processed Visium objects were obtained from the Human Cell Atlas (project e9f36305-d857-44a3-93f0-df4e6007dc97), with clinical metadata from Kuppe et al. [48]. The dataset contained 91,517 spots across 28 sections from 20 patients. We used the nine published computational cell-type-niche labels and retained the author-defined tissue categories: 14 myogenic-enriched, nine ischaemic-enriched, and five fibrotic-enriched sections. These categories describe tissue histomorphology; the myogenic-enriched category includes control tissue and remote or border regions from infarcted hearts. Array positions were transformed to an equilateral hexagonal coordinate system, and local and sample-level BD were calculated separately for each section using *k* = 6.

Relative BD was the primary sample-level summary. Sections in which one niche occupied more than 80% of spots were flagged, and group medians were also calculated after excluding these sections. Comparisons were descriptive, with each section contributing equally to group summaries and repeated patient identities indicated in the plots.

Neighborhood-size sensitivity was assessed using the same published niche labels and all 28 retained sections, as described in Supplementary Methods and Supplementary Figure S11.

#### 4.4.3 Miyoshi frontal-cortex Visium analyses

The author-processed Miyoshi frontal-cortex object was obtained from GEO accession GSE233208 and contained 115,451 spots across 39 sections [49]. The cohort comprised 10 cognitively normal controls, nine early-stage Alzheimer’s disease (AD), 10 late-stage AD, and 10 Down-syndrome-associated AD samples. We analyzed the published BayesSpace *q* = 9 annotation, comprising six cortical and three white-matter regions, and a second label definition obtained by merging WM1–WM3 into a single white-matter label while retaining the six cortical labels. Both definitions used the same retained spots, spot order, transformed Visium coordinates, and directed *k* = 6 nearest-neighbor search settings within each section, excluding the focal spot. We compared local raw BD maps and sample-level raw BD, expected raw BD, and relative BD values, with the analytic expectation calculated from the label counts for each definition. Sensitivity to label granularity was assessed using paired changes in raw BD, expected raw BD, and relative BD and the Spearman correlation of section-level relative BD between the two definitions. The representative section was selected as the 20th of 39 sections ordered by increasing relative BD under the original annotation.

Neighborhood-size sensitivity was assessed using the original nine-region *q* = 9 annotation and all 39 retained sections, as described in Supplementary Methods and Supplementary Figure S11.

#### 4.4.4 Maynard DLPFC Visium analyses

The Maynard dorsolateral prefrontal cortex (DLPFC) data were obtained from the `spatialLIBD`Bio-conductor package [50, 53]. We analyzed 12 sections from three donors, retaining 47,329 spots with published manual cortical-layer (L1–L6) and white-matter annotations after excluding 352 unlabeled spots. Local and sample-level BD were calculated separately for each section using *k* = 6. Six directly adjacent section pairs were identified using the deposited donor, within-block position, and replicate fields, and absolute differences in relative BD within pairs were summarized descriptively.

#### 4.4.5 de Oliveira colorectal-cancer Visium HD and Xenium analyses

P2CRC Visium HD and Xenium data from de Oliveira et al. [51] were obtained through the `OSTA.data`resource [5]. The analyses included 545,850 nonempty 8 *µ*m Visium HD bins and 339,681 nonempty Xenium cells from one section per platform. For Visium HD, 1,000 variable genes were selected before smoothing; Xenium analyses used all 422 imported genes. Expression was normalized and smoothed over *k*_*s*_ = 18 neighbors across all retained locations, followed by log transformation. For each platform, PCA with 15 components and *k*-means clustering with *K* = 15 were fitted on a randomly selected 30,000-location subset; all locations were then projected into the fitted PC space and assigned to the nearest cluster centroid. Local and sample-level BD were calculated using *k* = 8 for Visium HD and *k* = 6 for Xenium. BD was also calculated for the 15 Visium HD labels distributed with the source object. Feature-selection and fitting subsets, normalization, and full-data projection are detailed in Supplementary Methods.

### 4.5 Simulation designs

The composition simulation used 900 locations on a 30 × 30 grid and a directed four-nearest-neighbor graph. We generated 75 random arrangements at each exact 10:90, 30:70, and 50:50 two-label composition. Five additional maps with 450 locations per label compared coherent domains, partial label exchange, random placement, and a checkerboard. Three further maps contained contiguous domains at the same three compositions while retaining the same graph, interface orientation, and interface extent. Raw, expected, and relative BD were calculated for every map, with expectations obtained analytically from the label counts.

The over-smoothing simulation used 3,600 locations on a 60 × 60 grid with two broad domains and an 81-location circular island. We compared no smoothing with uniform bandwidths from 2.5% to 30% of the coordinate range, holding the selected feature set, number of principal components (10), and number of clusters (*K* = 3) fixed. PCA was recomputed for each expression matrix. Domain recovery was assessed using ARI and island F1, and raw and relative BD were calculated using a directed four-nearest-neighbor graph. This analysis used one illustrative count realization. Count-generation procedures, label-exchange rules, complete simulation settings, and recovery-score definitions are provided in Supplementary Methods.x

## Code and data availability

The smoothclust R package is available from Bioconductor at https://bioconductor.org/packages/smoothclust, and its source code is maintained at https://github.com/lmweber/smoothclust. Scripts to reproduce the preprocessing, analyses, figures, and runtime evaluations are available at https://github.com/lmweber/smoothclust-analyses. All datasets analyzed are publicly available from the repositories and accessions specified in the Methods and Supplementary Table S1.

## Competing interests

The authors declare that they have no competing interests.

## Funding

LMW was supported by the National Human Genome Research Institute of the National Institutes of Health (R00HG012229).

## Acknowledgments

We thank colleagues at Johns Hopkins Bloomberg School of Public Health and the Lieber Institute for Brain Development for valuable early discussions that helped motivate the development of the methods and analyses presented here.

## Supplementary Materials

### Supplementary Methods

#### Smoothing weights and spatial scale

Let ***r***_*i*_ be row *i* of *S, d*_*ij*_ = ∥***r***_*i*_ − ***r***_*j*_∥_2_, and *h* = *bR*, where *R* is the larger horizontal or vertical coordinate range. Uniform-radius smoothing uses

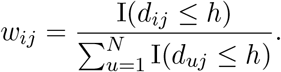

For exponential-kernel smoothing, with *h >* 0,

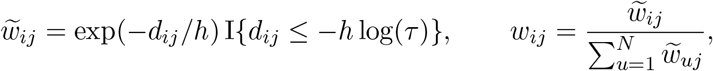

where unnormalized weights below *τ* = 0.05 are omitted by default. For nearest-neighbor smoothing, let ℳ_*j*_ contain the *k*_*s*_ nearest locations other than *j*. Then

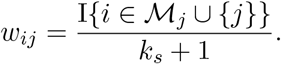

All three smoothers include the focal location. The default *k*_*s*_ = 18 corresponds to two neighbor layers in a complete Visium hexagonal layout. Equal nominal settings do not imply equal physical scales: *bR* is a hard radius for uniform smoothing but an exponential length scale for the kernel, whose retained support extends to approximately 3*bR* at *τ* = 0.05. Nearest-neighbor smoothing fixes neighbor count; smoothing scale is distinct from the neighborhood used to calculate BD.

#### Input handling and normalization

Expression columns, coordinate rows, and labels were supplied in matching order, with tissue sections analyzed separately. Size factors were supplied explicitly or calculated from column sums of the selected expression matrix, then rescaled to mean one. Dataset-specific normalization references are given below.

#### Derivation of the random-label expectation

For a fixed graph without self-relations on *N >* 1 locations, let *π* uniformly reassign labels while preserving counts *n*_1_, … , *n*_*L*_. Writing *p*_ℓ_ = *n*_ℓ_*/N* , any pair *i* ≠ *j* satisfies

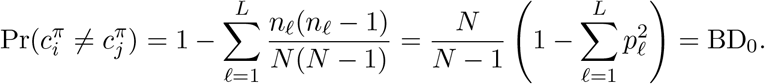

The factor *N/*(*N* − 1) accounts for sampling labels without replacement. Linearity of expectation, without requiring independent edge indicators, gives

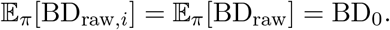

Consequently, E_*π*_[rBD] = 1 whenever BD_0_ *>* 0. The same expectation holds for fixed nonnegative weights on distinct location pairs normalized by their positive total. Observed BD and its random-label variability remain graph-dependent.

Merging labels with counts *n*_*a*_ and *n*_*b*_, while retaining the same locations, changes the expectation by

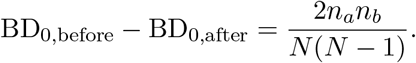

The expectation decreases when both labels are present; repeated application gives the result for merging multiple labels. On an unchanged graph, neither raw BD nor its expectation can increase after merging, but relative BD can increase if its denominator decreases proportionally more than its numerator.

#### Relationship to SHouT graph homophily and focal-label agreement

For a directed *k*-nearest-neighbor graph, global edge homophily satisfies

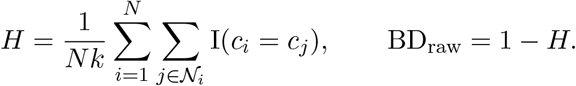

Equality with the complement of SHouT’s global homophily requires matching neighbor relations, multiplicities, and weights, but not graph symmetry [43]. The fraction of neighbors sharing the focal label is 1 −BD_raw,*i*_ when the focal location is excluded. If egophily *e*_*i*_ includes the focal location and the same *d*_*i*_ neighbors, then 1 − *e*_*i*_ = {*d*_*i*_*/*(*d*_*i*_ + 1)}BD_raw,*i*_. SHouT’s local homophily instead summarizes edges within a neighborhood and is distinct from this focal-label comparison.

#### Relationship to Geary’s *C*

For the directed *k*-nearest-neighbor graph, let *a*_*ij*_ = I(*j* ∈ N_*i*_), *a*_*ii*_ = 0, and *A*_0_ = ∑_*i,j*_ *a*_*ij*_ = *Nk*. For nonconstant *x*, Geary’s statistic is [39]

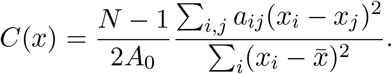

For at least two observed labels, define *z*_*i*ℓ_ = I(*c*_*i*_ = ℓ) and *v*_ℓ_ = ∑_*i*_(*z*_*i*ℓ_ − *p*_ℓ_)^2^ = *Np*_ℓ_(1 − *p*_ℓ_). Since 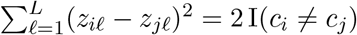,

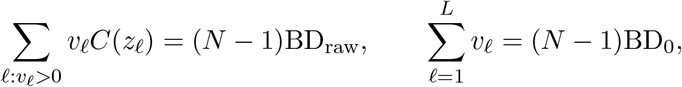

and therefore

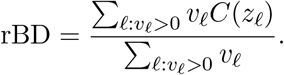

Thus, relative BD is a variance-weighted, not generally unweighted, mean of indicator-specific Geary statistics. With two labels, the complementary indicators have equal Geary statistics, giving rBD = *C*(*x*) for either binary indicator *x*. These identities require matching locations and spatial weights. For a variable-degree graph with all *d*_*i*_ *>* 0, the corresponding mean of local fractions uses row-normalized weights *a*_*ij*_ = I(*j* ∈ *N*_*i*_)*/d*_*i*_, with *A*_0_ = *N* .

##### Relationship to the initial stEDGE boundary signal

stEDGE constructs an initial boundary signal from consensus co-assignment across labelings [46]. For *M* labelings of the same locations, define

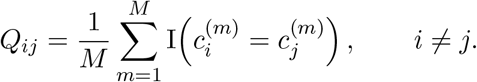

For a common neighborhood N_*i*_ excluding *i*, its complement-of-consensus score is

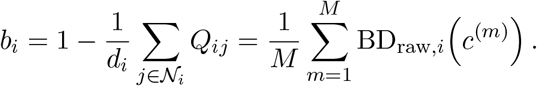

This identity requires matching neighborhoods, normalization, and exclusion of self-relations. It concerns only the initial boundary signal, not stEDGE’s subsequent domain reconstruction, hierarchical organization, or transition index, and does not imply equivalence to relative BD.

#### GBM preprocessing and feature selection

GEO matrices were restricted to in-tissue spots with at least 1,000 total UMIs, at most 20% mitochondrial UMIs, finite totals and mitochondrial percentages, and positive nonmitochondrial totals. Genes with zero counts across retained spots were removed; the shared analysis set retained nonmitochondrial genes with at least 100 counts in at least one section, excluding spots with zero counts after filtering. Both workflows used size factors based on all nonmitochondrial genes before low-expression filtering, rescaled to mean one within sections, and pseudocount one. Features were selected separately from smoothed and unsmoothed log-expression using `scran::modelGeneVar()`and `scran::getTopHVGs()`with `prop=0.1`and otherwise default arguments; fitted scores ranked features for smoothed expression. PCA was fitted separately with `scater::runPCA()`, the selected genes, 50 components, and otherwise default settings.

#### External clustering settings

BayesSpace 1.22.0 used `spatialPreprocess()`with ⌈0.1*G*⌈ variable genes from the *G* shared input genes, 15 principal components, and log normalization with the above per-section size factors. Clustering used *q* = 5, *k*-means initialization, the *t* model with equal precision, 50,000 MCMC iterations, 1,000 burn-in iterations, and thinning interval 100 [9]. BANKSY (`Banksy`1.8.1) and ClustSIGNAL 1.4.1 used `scuttle::logNormCounts()`with the same size factors and pseudocount one, followed by the above feature-selection procedure (`prop=0.1`). BANKSY used *M* = 0, *λ* = 0.8, *k*_geom_ = 15, no azimuthal Gabor features, 50 principal components, and *K* = 5 *k*-means [17]. ClustSIGNAL used default labeling settings through `clustSIGNAL(spe cs, samples=“sample id”, threads=14, outputs=“s”)`[18]. Fourteen threads were requested where supported; effective parallelism could vary by stage. Seeds were deterministic by section.

#### Greenwald annotation matching and summaries

Annotations in `visium metadata.csv`were matched by section identifier and spot barcode after mapping processed section names to deposited identifiers. Unmatched spots received missing annotations. Deposited labels, including layer annotations for Chromatin-Reg-dominant spots, were retained. Each annotation field used a graph of spots with nonmissing labels. Structured-zone fraction was the proportion of `org1`-labeled spots assigned `struct`;

MES-Hyp fraction was the proportion assigned `MES.Hyp`among spots with nonmissing `mp`within the `org1`-labeled subset. Metaprogram composition used all spots with nonmissing `mp`per section. Regional comparisons included 12 sections from ZH1007, ZH1019, ZH881, and ZH916. GBM graphs used `pxl col in fullres`and `pxl row in fullres`in full-resolution image pixels.

#### GBM smoothing and graph-scale sensitivity

Uniform and kernel smoothing at *b* = 0.05 were compared with nearest-neighbor smoothing at *k*_*s*_ = 18, without matching effective spatial reach. Parameter sensitivity used *b* = 0.010, 0.025, 0.050, 0.075, 0.100, 0.150 and *k*_*s*_ = 2, 6, 12, 18, 30, 42, respectively. BD neighborhood sensitivity held labels fixed and used *k* = 2, 4, 6, 8, 12, 18, 24, 36. All settings were evaluated independently by section for *K* = 3, … , 8.

#### Kuppe and Miyoshi annotation matching and coordinates

Kuppe metadata, expression columns, and image coordinates required identical barcode order; deposited `celltype niche`labels were retained. Miyoshi coordinates were matched to spot identifiers in metadata order, using labels from the deposited `annotation`field. For both datasets, array coordinates were transformed as *x* = `col`*/*2 and 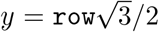. After excluding missing or empty labels, section-specific directed Euclidean graphs used *k* = 6, excluding focal locations, with `BiocNeighbors::findKNN()`, four requested threads, and otherwise default search settings.

#### Neighborhood-size sensitivity for published annotations

For the Kuppe nine-niche labels (28 sections) and Miyoshi nine-region BayesSpace annotation (39 sections), BD at *k* = 18 and *k* = 36 was compared with *k* = 6 (Supplementary Figure S11). Directed Euclidean graphs were rebuilt per section and neighborhood size, excluding focal locations while retaining the same spots, coordinates, labels, spot order, and search settings. Recomputed *k* = 6 values were checked against the saved baseline, and analytic expectations were verified unchanged. For raw and relative BD, summaries comprised Spearman correlations with *k* = 6, signed changes (larger *k* minus *k* = 6), and median and maximum absolute changes. Kuppe category medians and interquartile ranges used all sections with equal section weighting at each *k*; comparisons were descriptive, not treating repeated patient sections as independent replicates.

#### Maynard annotations

The processed `sce`object distributed through `spatialLIBD`(version 1.24.0) was used, as described in Supplementary Table S1. Labels were from `layer guess reordered short`. All nonmissing, nonempty labels were retained, including 534 spots marked `discard=TRUE`; 352 unannotated spots were excluded. Adjacent pairs used `subject`, `position`, `replicate`, and `subject position`, pairing replicates 1 and 2 within donor and within-block position.

#### High-resolution preprocessing and sketching

Visium HD and Xenium locations with positive imported count totals were retained; these full-library totals provided mean-one size factors. Visium HD genes were eligible if detected in at least 0.1% of retained bins, excluding mitochondrial genes. The 1,000 genes with highest variance of log_2_(1 + normalized counts) were selected on a random 25,000-bin subset, with library totals rescaled to mean one within that subset. Bins with zero counts among selected genes were retained. Xenium used all 422 imported genes. Normalization, nearest-neighbor smoothing, and log transformation used all retained locations. PCA and clustering were fitted on a separate random 30,000-location subset for each platform; the Visium HD fitting subset was sampled separately from its feature-selection subset. All subsets were sampled without replacement. PCA used `irlba::prcomp irlba()`, centering without scaling; all locations were projected using the fitted centering and loadings. *k*-means used three starts and at most 500 iterations, followed by nearest-centroid assignment of all locations. Visium HD coordinates were `array col`and `array row`(one unit = 8 *µ*m); Xenium used the imported spatial coordinates.

#### Composition simulation

The composition simulation used a 30 × 30 grid with exact A:B label counts of 90:810, 270:630, or 450:450. We generated 75 random label arrangements per composition. For the fixed 450:450 composition, a coherent map assigned A to *x* ≤ 15 and B otherwise. To introduce partial mixing, we randomly exchanged labels at 10% or 30% of locations while preserving the label counts. We also generated a random arrangement and a checkerboard based on the parity of *x* + *y*. Additional straight-interface maps assigned A to *x* ≤ *t*, with *t* = 3, 9, 15, and B otherwise, varying composition while retaining the same interface orientation and extent. All calculations used directed four-nearest-neighbor graphs without periodic boundaries.

#### Over-smoothing simulation

The over-smoothing simulation used 3,600 locations on a 60 × 60 grid with two broad domains and a small circular third-domain island containing 81 locations. For 120 genes, baseline means were drawn from a gamma distribution (shape and rate both 1.5), with 15 distinct marker genes per domain having twofold higher means. Negative-binomial counts (size parameter 1) were generated with location-specific log-normal library multipliers (log-standard deviation 0.35), rescaled to mean one. Counts were normalized using observed library totals. The 75 most variable genes from unsmoothed log-transformed normalized expression were selected and held fixed across settings. We compared an unsmoothed baseline with eight uniform-smoothing bandwidths ranging from 2.5% to 30% of the coordinate range. For each setting, PCA was recomputed, followed by *K* = 3 *k*-means clustering using the first 10 PCs. Domain recovery was assessed using the adjusted Rand index (ARI) and island F1, defined for true island *I* and inferred clusters *C*_*j*_ as

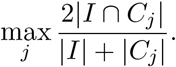

Counts, normalization factors, selected features, coordinates, and directed four-nearest-neighbor settings were held fixed across bandwidths. Cluster labels were aligned to the true labels only for visualization. This analysis used one illustrative count realization.

## Supplementary Tables

**Supplementary Table S1:** Spatial transcriptomics datasets and analysis inputs used in this study.

| Dataset | Reference and data source | Platform | Samples analyzed | Spatial locations | Expression inputs and labels |
| --- | --- | --- | --- | --- | --- |
| GBM/glioma | Greenwald et al. (2024) [47]; GEO GSE237183 | Visium, fresh frozen | 19 sections<sup>a</sup> | 34,699 retained spots | 14,301 shared input genes; smoothclust, baseline, and external-method labels; deposited **org1**, **org2**, **mp**, **layer** |
| Myocardial infarction | Kuppe et al. (2022) [48]; HCA<sup>b</sup> | Visium, fresh frozen | 28 sections from 20 patients | 91,517 annotated spots | Nine published cell-type-niche labels |
| Human frontal cortex | Miyoshi et al. (2024) [49]; GEO GSE233208<sup>c</sup> | Visium, fresh frozen | 39 sections | 115,451 deposited spots | Published nine-region BayesSpace annotation (six cortical, three white-matter regions) and WM1–WM3-merged definition |
| Human DLPFC | Maynard et al. (2021) [50]; **spatialLIBD** [53] archive<sup>d</sup> | Visium, fresh frozen | 12 sections from 3 donors; 6 adjacent pairs | 47,329 annotated spots | Published manual L1–L6 and white-matter labels |
| Visium HD CRC | de Oliveira et al. (2025) [51]; **OSTA.data**<sup>e</sup> | Visium HD, FFPE | One P2CRC section | 545,850 nonempty 8 μm bins | 1,000 selected variable genes; 15 smoothclust-derived domains; 15 distributed cluster labels |
| Xenium CRC | de Oliveira et al. (2025) [51]; **OSTA.data**<sup>e</sup> | Xenium In Situ, FFPE | One P2CRC section | 339,681 nonempty segmented cells | All 422 imported genes; 15 smoothclust-derived domains |

All platforms are from 10x Genomics. <sup>a</sup>The GBM/glioma clustering cohort comprised 13 GBM and six IDH-mutant glioma sections. Published annotations were available for 13 sections; regional analyses used 12 sections from four tumors. Annotation file: **Inputs/general/visium\_metadata.csv**. <sup>b</sup>HCA project e9f36305-d857-44a3-93f0-df4e6007dc97; 28 sections represent 27 patient-region samples. <sup>c</sup>Processed object: **GSE233208\_Human\_visium\_ADDS\_seurat\_processed.rds.gz**. <sup>d</sup>Archived processed object: **Human\_DLPFC\_Visium\_processedData\_sce\_scran\_spatialLIBD.Rdata**; annotation retention and software version are described in Supplementary Methods. <sup>e</sup>Serial sections from the same P2CRC FFPE block. **OSTA.data** resources: **VisiumHD\_HumanColon\_Oliveira** and **Xenium\_HumanColon\_Oliveira**. Visium HD inputs: **binned\_outputs/square\_008um/filtered\_feature\_bc\_matrix.h5** and distributed labels in **clustering.csv.gz**; Xenium count input: **cell\_feature\_matrix.h5**.

**Supplementary Table S2:** Runtime for representative core smoothclust workflows.

| Workflow | Sections | Locations | Input genes | Feature-reduction and clustering scope | Requested<br>NN<br>threads | Runtime |
| --- | --- | --- | --- | --- | --- | --- |
| GBM Visium | 19 | 34,699 | 14,301 | Independent full-data workflow per section, $K = 5$ | 14 | 33.9 s |
| P2CRC Visium HD | 1 | 545,850 | 1,000 | 30,000-bin PCA/clustering fit; full-data projection and assignment, $K = 15$ | 8 | 54.0 s |
| P2CRC Xenium | 1 | 339,681 | 422 | 30,000-cell PCA/clustering fit; full-data projection and assignment, $K = 15$ | 8 | 16.2 s |

Timings were measured on a 2023 MacBook Pro (Apple M3 Max, 14 cores, 36 GB memory). NN denotes nearest-neighbor searches; thread counts are requested values, and effective parallelism may vary. Core runtimes include smoothing, feature reduction, clustering or full-data assignment, and BD; they exclude data import, neighbor-distance diagnostics, file output, plotting, and the additional BD calculation for distributed Visium HD labels. GBM variable-gene selection and PCA were timed; Visium HD variable-gene selection occurred before timing, and Xenium used all 422 imported genes. The GBM $K = 5$ runtime was reconstructed from component timers in the $K = 3, \dots, 8$ analysis; Visium HD and Xenium were each timed once.

## Supplementary Figures

**Supplementary Figure S1:**
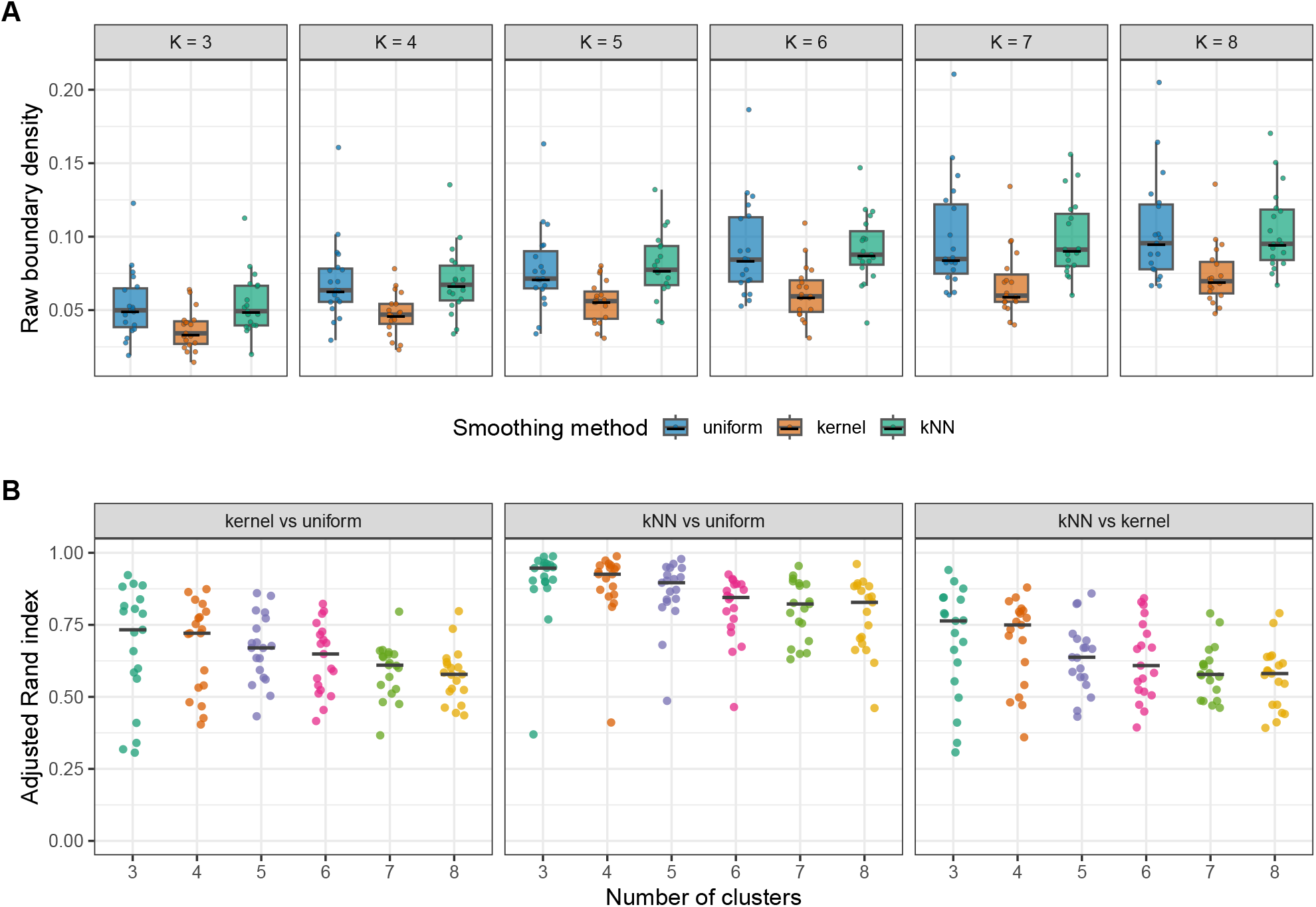
GBM smoothing method sensitivity analysis. (A) Raw BD for smoothclust labels generated using uniform-radius, kernel, and *k*-nearest-neighbor smoothing across *K* = 3, … , 8. Boxes show medians and interquartile ranges, with whiskers extending to observations within 1.5 interquartile ranges; points represent individual GBM/glioma tissue sections and horizontal bars indicate medians. (B) Pairwise adjusted Rand index (ARI) between labelings from the three smoothing methods, grouped by cluster number and faceted by smoothing-method pair. Points represent individual sections and bars indicate medians. The BD metric was calculated using *k* = 6; uniform and kernel smoothing used *b* = 0.05, and *k*-nearest-neighbor smoothing used *k*_*s*_ = 18. In these settings, *h* = *bR* is a hard radius for uniform smoothing and an exponential length scale with wider truncated support for kernel smoothing, whereas the *k*-nearest-neighbor setting fixes neighbor count.

**Supplementary Figure S2:**
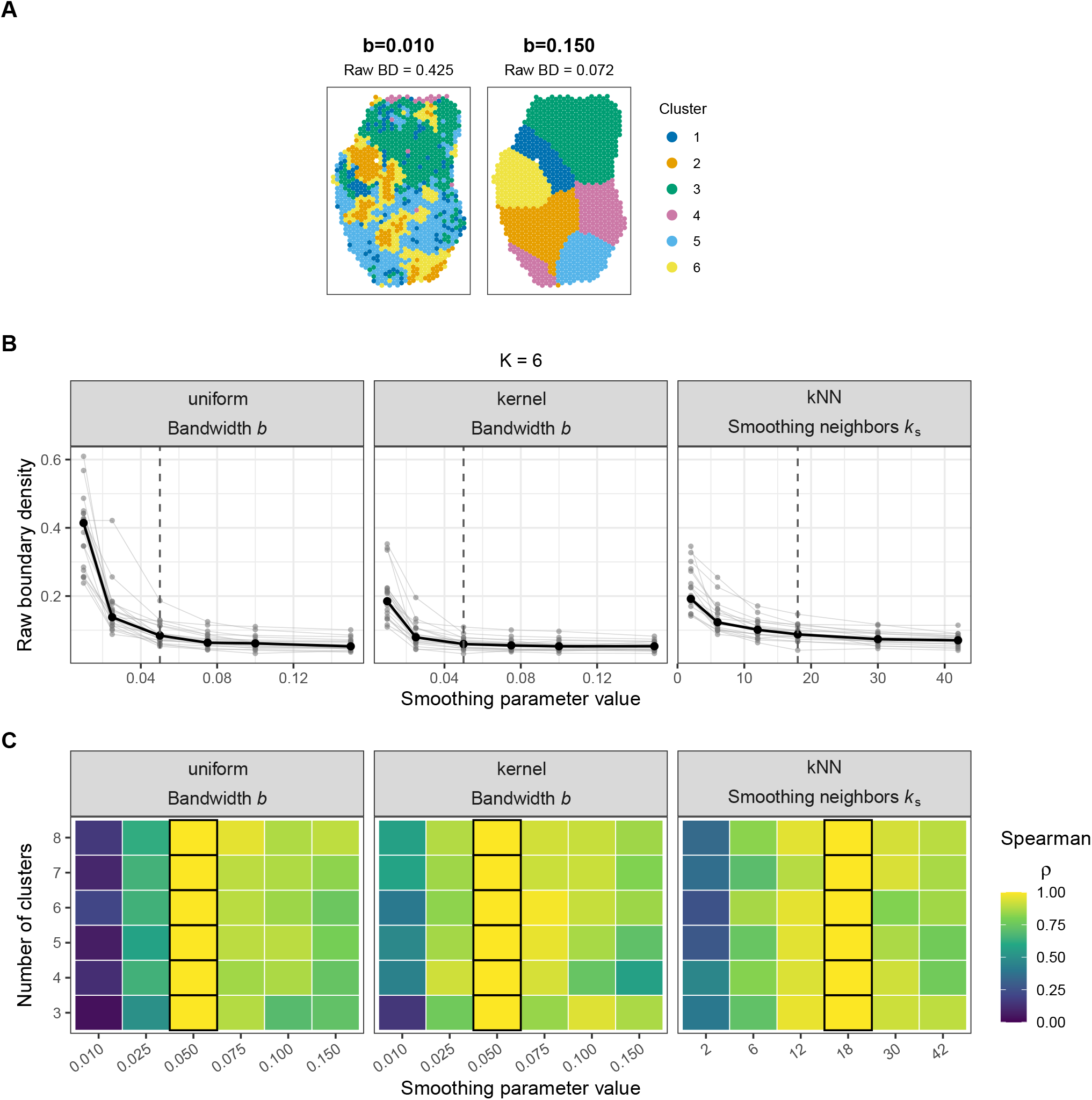
GBM smoothing parameter sensitivity analysis. (A) Representative domain maps for section zh1019t1 with *K* = 6 using uniform smoothing at smaller bandwidth *b* = 0.010 (raw BD = 0.425; left) and larger bandwidth *b* = 0.150 (raw BD = 0.072; right). (B) Raw BD as a function of smoothing bandwidth or smoothing-neighbor count at *K* = 6. Gray lines represent sections and black points/lines indicate medians. Dashed vertical lines mark defaults. Uniform-smoothing bandwidth is expressed relative to each section’s larger post-filtering coordinate range. (C) Spearman rank correlation between sample-level raw BD values at each setting and the corresponding default across *K* = 3, … , 8. Black outlines mark default settings, whose correlations equal 1 by self-comparison. Defaults were *b* = 0.05 for uniform and kernel smoothing and *k*_*s*_ = 18 for nearest-neighbor smoothing. The BD metric was calculated using *k* = 6.

**Supplementary Figure S3:**
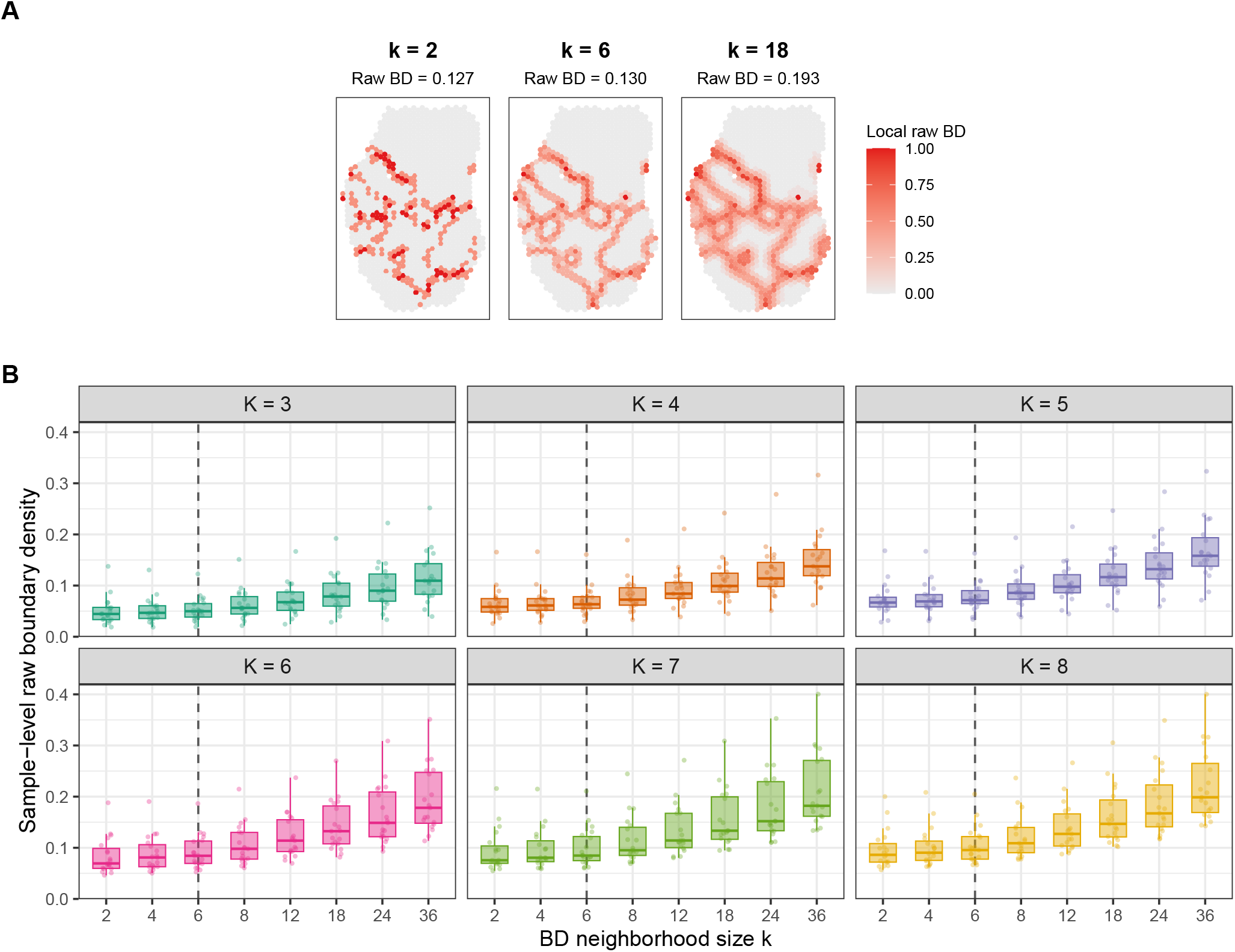
BD metric neighborhood-size sensitivity in the GBM cohort. (A) Representative local raw BD maps for section zh1019t1 with *K* = 6, calculated using small (*k* = 2), default (*k* = 6), and large (*k* = 18) neighborhoods, from left to right. Values below each heading give sample-level raw BD. (B) Sample-level raw BD distributions across neighborhood sizes *k* = 2, 4, 6, 8, 12, 18, 24, 36, faceted by cluster number *K* = 3, … , 8. Points represent the 19 sections; boxes show medians and interquartile ranges. Dashed vertical lines mark the default *k* = 6. smoothclust labels were held fixed.

**Supplementary Figure S4:**
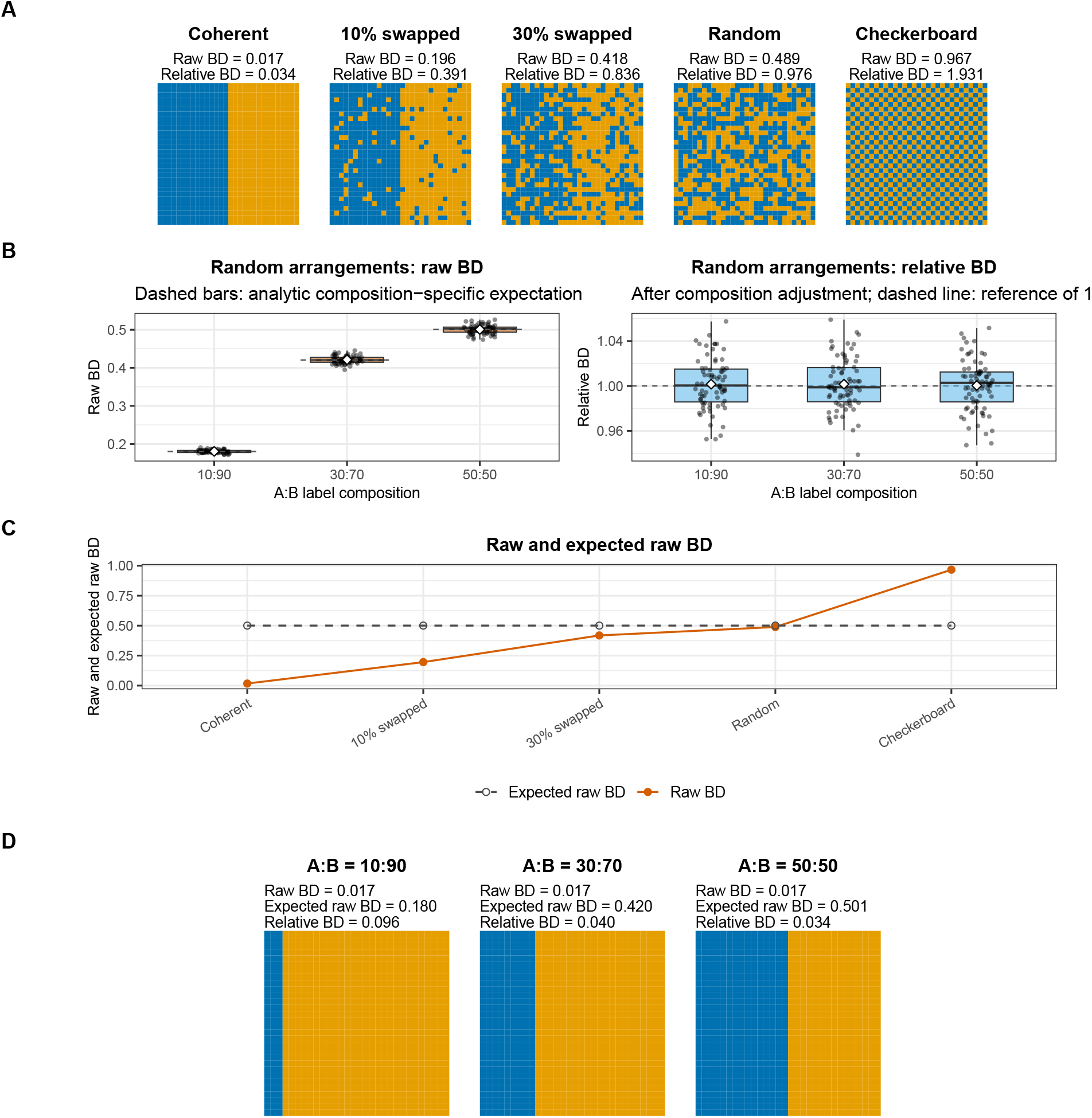
Illustrating the composition adjustment underlying relative BD. (A) Two-label configurations on a 30 × 30 regular grid, each with exactly 450 locations per label. In panels A and D, label A is blue and label B is orange. (B) Raw and relative BD across 75 independent random arrangements at each exact label composition. Dashed bars mark analytically expected raw BD, and the dashed line marks the relative-BD reference of 1. Diamonds mark means. (C) Observed and analytically expected raw BD for the fixed-450:450 organization series. The random map is one realization; the checkerboard deliberately contains more label transitions than the random-label expectation. At fixed composition, raw and relative BD differ by a constant scaling; relative BD values are also shown in the map subtitles in panel A. (D) Straight-interface maps with 10:90, 30:70, and 50:50 compositions on the same grid, with raw BD, expected raw BD, and relative BD shown for each. The graph, interface orientation, and interface extent are fixed across these three maps, illustrating how approximately equal observed transition frequencies can yield different relative BD because the composition-specific references differ. All BD calculations used directed four-nearest-neighbor graphs; expected raw BD was calculated analytically.

**Supplementary Figure S5:**
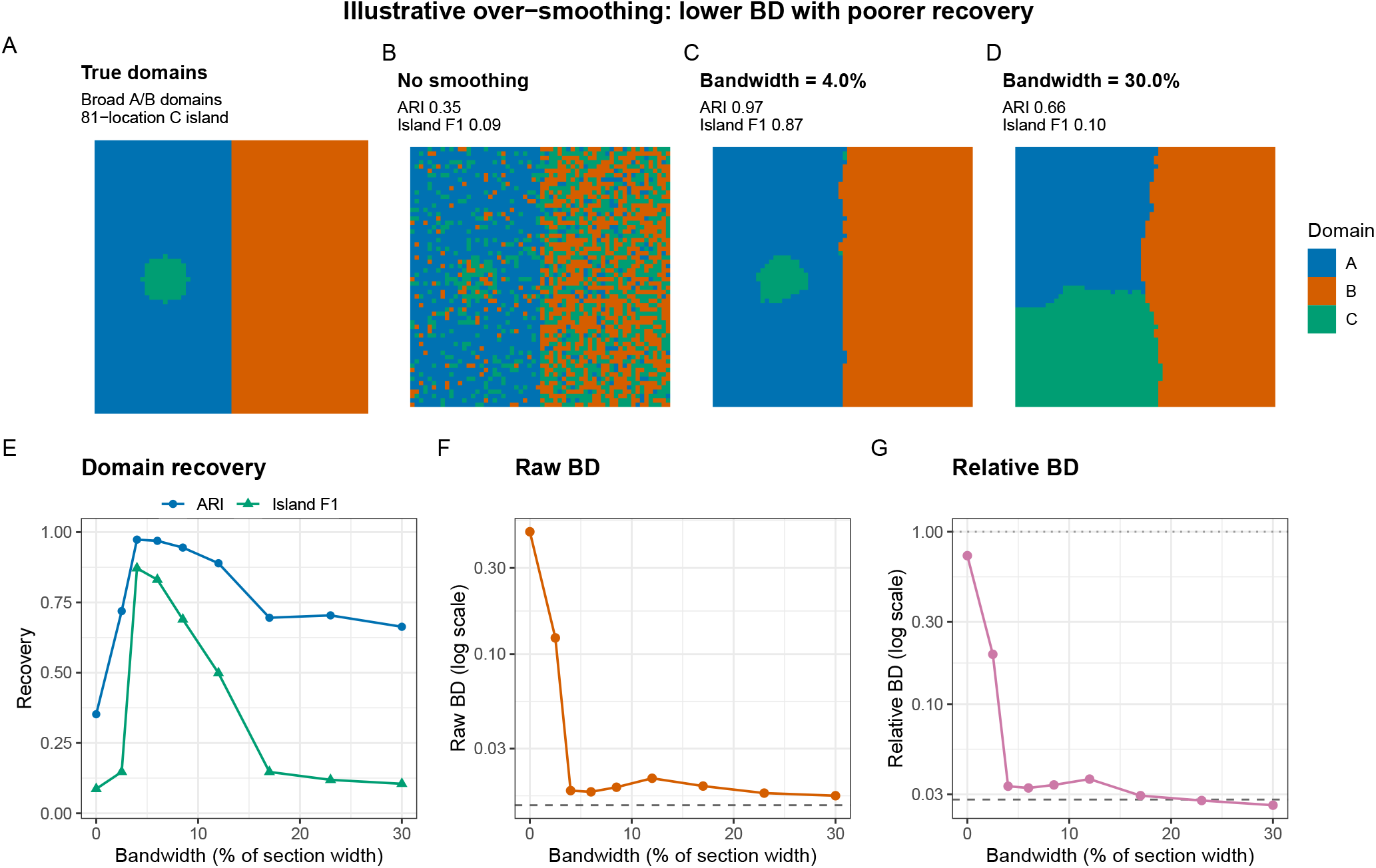
Illustrative over-smoothing simulation with lower BD metric values and poorer domain recovery. (A) True labels for a three-domain count simulation on a 60 × 60 grid with broad left and right domains and an 81-location circular third-domain island. (B–D) Inferred three-cluster maps without smoothing, at the 4% uniform bandwidth (highest ARI and island F1 in this realization), and at the largest evaluated bandwidth of 30% of the coordinate range. Cluster identities were aligned to truth only for display. (E) ARI and island F1 across all evaluated bandwidths. (F) Raw BD and (G) relative BD, the composition-adjusted summary, for the inferred labels; dashed lines mark the values for the true labels and the dotted line in panel G marks the fixed-count random-label reference. Both F and G use logarithmic y-axes. The feature set, number of principal components (10), *K* = 3, coordinates, and directed four-nearest-neighbor settings were held fixed; PCA was recomputed for each expression matrix. This analysis used one illustrative count realization.

**Supplementary Figure S6:**
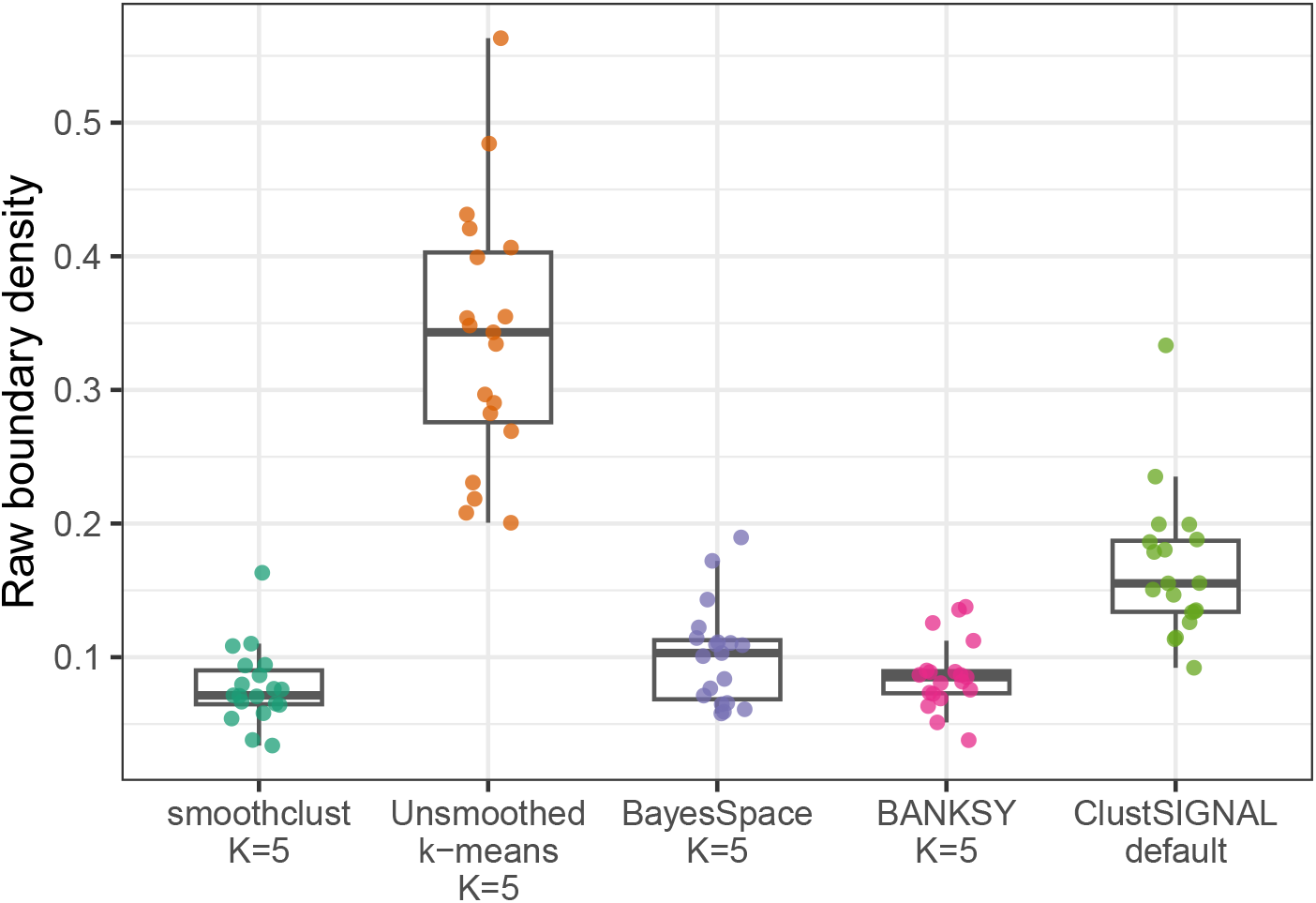
Sample-level raw BD for labels generated by different clustering methods. Raw BD for smoothclust, the matched unsmoothed *k*-means baseline, and the external clustering algorithms BayesSpace, BANKSY, and ClustSIGNAL across 19 GBM/glioma sections. Boxes show medians and interquartile ranges, with whiskers extending to observations within 1.5 interquartile ranges; points represent individual sections. The first four methods are shown at *K* = 5; ClustSIGNAL used default parameters and returned 3–10 labels. All BD calculations used *k* = 6.

**Supplementary Figure S7:**
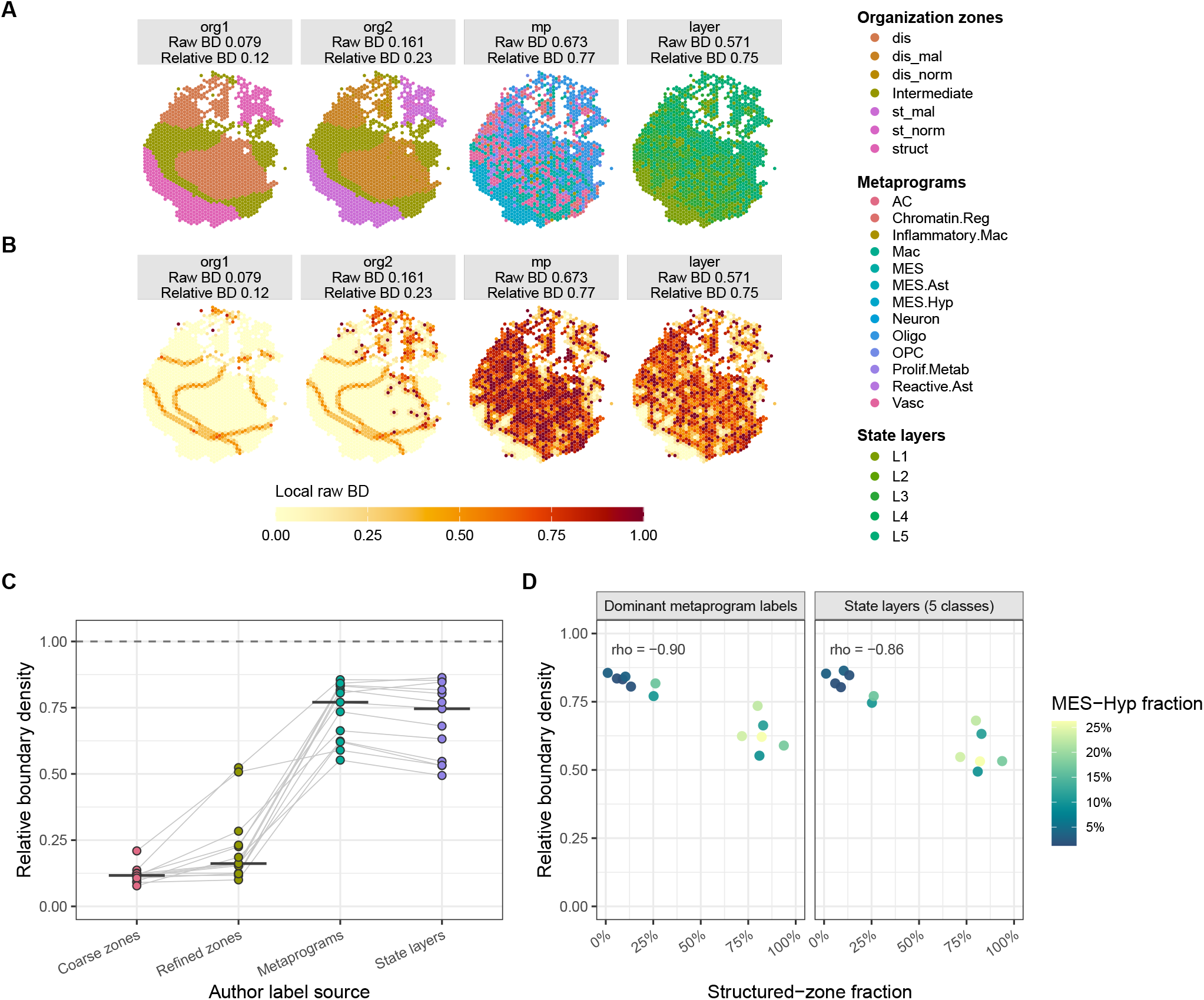
BD summarizes published GBM tissue annotations. (A) Representative Greenwald organization zones (`org1`and `org2`), dominant metaprograms (`mp`), and deposited state-layer annotations (`layer`) in section zh1019inf. Chromatin-Reg-dominant spots retained their deposited L1 assignments. (B) Local raw BD maps; facet labels report sample-level raw and relative BD. (C) Relative BD across the 13 annotated sections for Coarse zones (`org1`), Refined zones (`org2`), Metaprograms (`mp`), and State layers (`layer`); lines connect fields from the same section. Median raw BD was 0.049 for `org1`, 0.071 for `org2`, 0.673 for `mp`, and 0.571 for `layer`. The dashed line marks relative BD = 1. (D) Relative BD for `mp`and `layer`against structured-zone fraction; color indicates MES-Hyp fraction. Correlations are descriptive across 13 sections from five tumors. All BD calculations used *k* = 6.

**Supplementary Figure S8:**
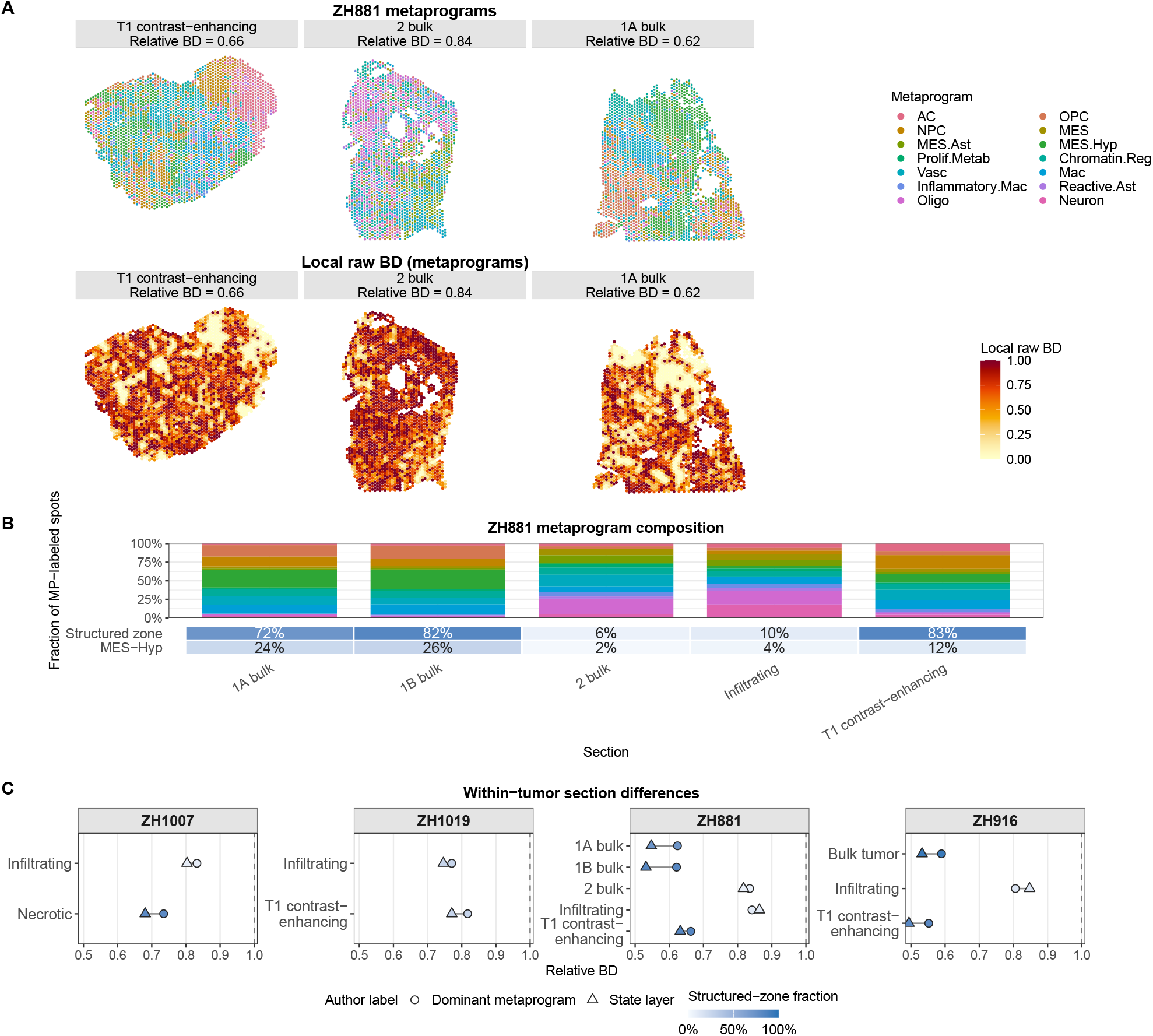
BD summarizes variation among regions within GBM tumors. (A) Dominant-metaprogram label maps and corresponding local raw BD maps for three ZH881 sections; headings report metaprogram relative BD. (B) Metaprogram composition across all five ZH881 sections, with structured-zone and MES-Hyp percentages below the composition bars. (C) Metaprogram and state-layer relative BD for 12 regional sections from four tumors; fill indicates structured-zone fraction and lines connect label sources from the same section. Dashed vertical lines mark relative BD = 1. State-layer values retain the deposited L1 assignments for Chromatin-Reg-dominant spots. BD was calculated on labeled-spot *k* = 6 graphs using the analytic fixed-count reference.

**Supplementary Figure S9:**
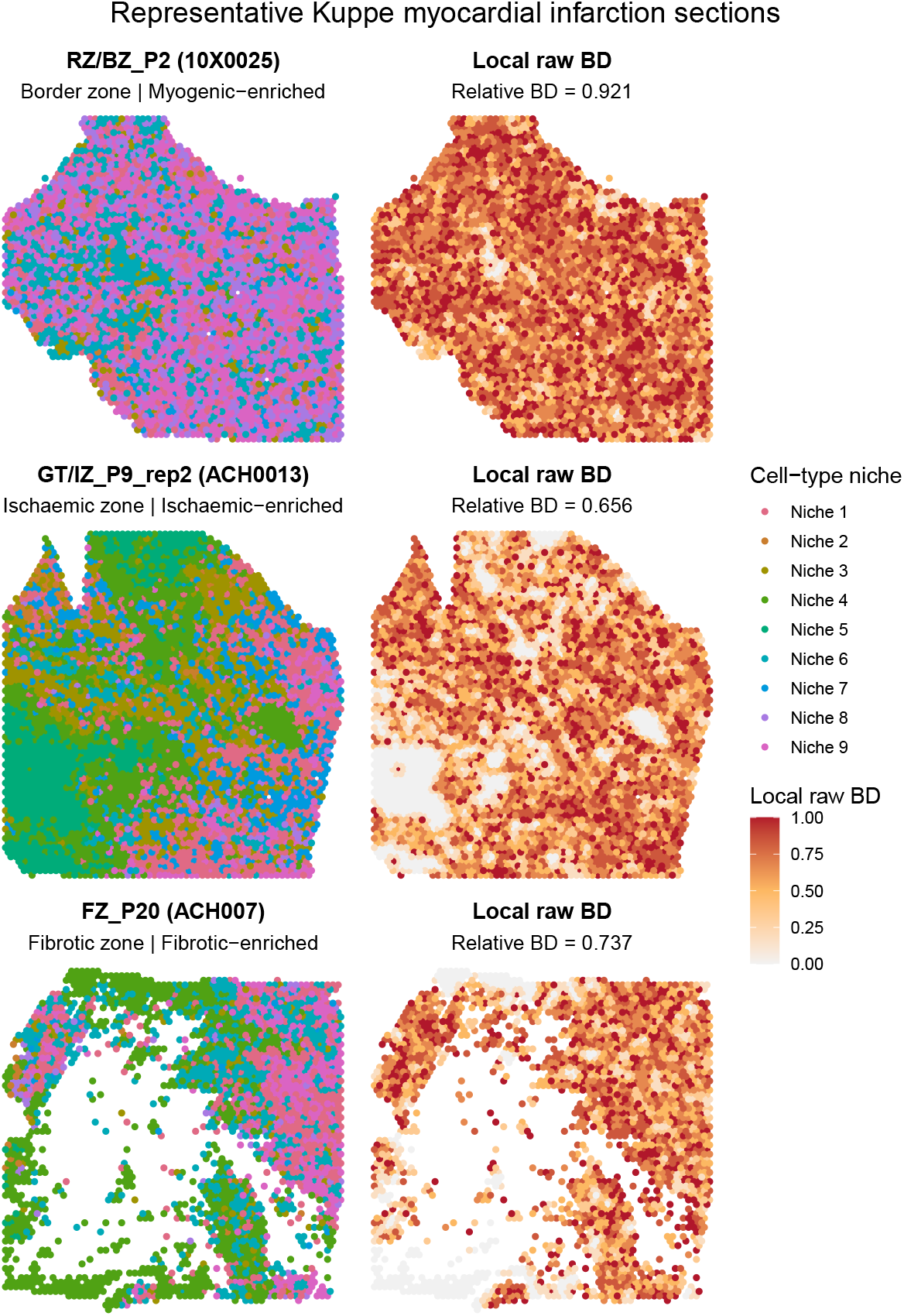
Published cell-type niches and local raw BD in representative myocardial sections. For each author-defined tissue category, we selected the section closest to its median relative BD: RZ/BZ P2 (myogenic-enriched; 0.921), GT/IZ P9 rep2 (ischaemic-enriched; 0.656), and FZ P20 (fibrotic-enriched; 0.737). Left panels show the nine published computational cell-type-niche labels; right panels show corresponding local raw BD, the fraction of each spot’s six nearest neighbors assigned to a different label.

**Supplementary Figure S10:**
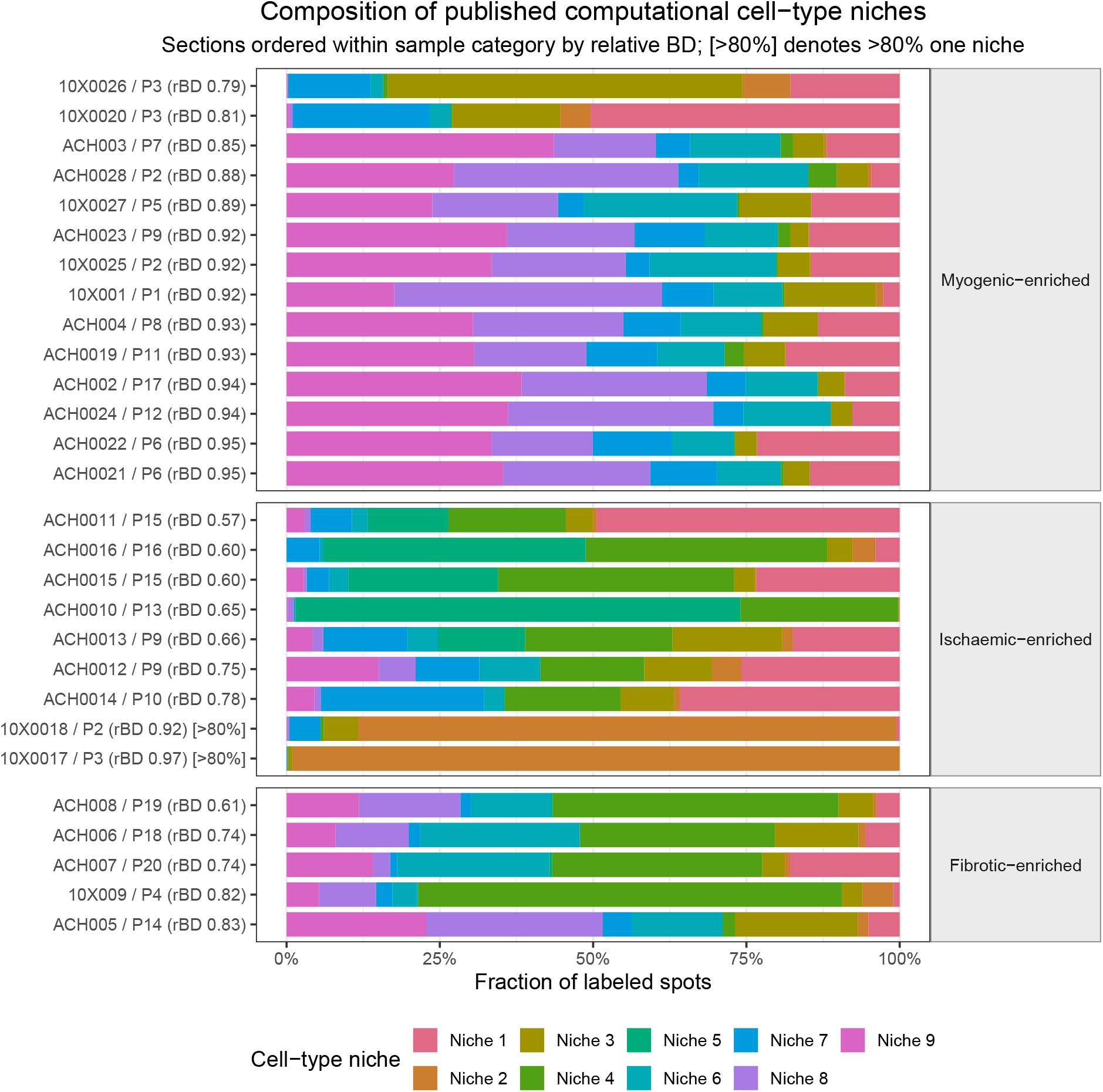
Composition of published myocardial cell-type niches across 28 sections. Fraction of spots assigned to each of the nine computational cell-type niches. Sections are grouped by the author-defined tissue category and ordered within each category by relative BD. Labels identify sections and patients and report relative BD; [*>* 80%] marks sections in which one niche occupies more than 80% of spots.

**Supplementary Figure S11:**
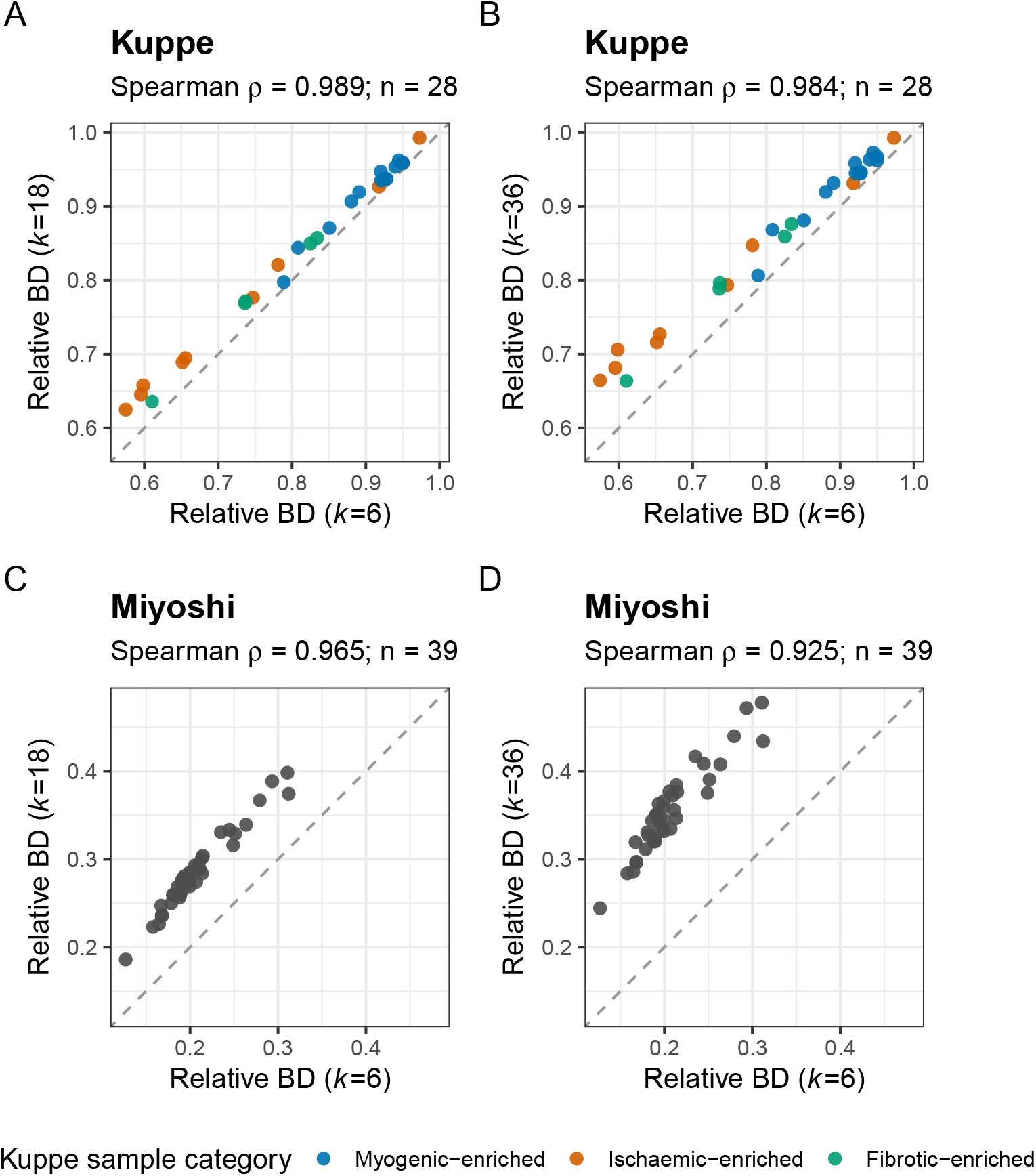
BD neighborhood-size sensitivity for published myocardial and cortical annotations. (A–B) Relative BD at *k* = 18 and *k* = 36, respectively, against *k* = 6 for the nine published Kuppe cell-type-niche labels across 28 sections from 20 patients; colors denote the author-defined tissue categories. (C–D) The corresponding comparisons for the original nine-region BayesSpace *q* = 9 annotation across 39 Miyoshi frontal-cortex sections. Each point is one section; dashed lines indicate equality. Axes have equal scaling, with common limits within each dataset’s row, and panels report Spearman correlations. For *k* = 18 and *k* = 36, respectively, median (maximum) absolute changes in relative BD were 0.025 (0.059) and 0.039 (0.108) for Kuppe, and 0.078 (0.096) and 0.145 (0.182) for Miyoshi. Section rankings remained strongly correlated while absolute values changed, particularly for Miyoshi.

**Supplementary Figure S12:**
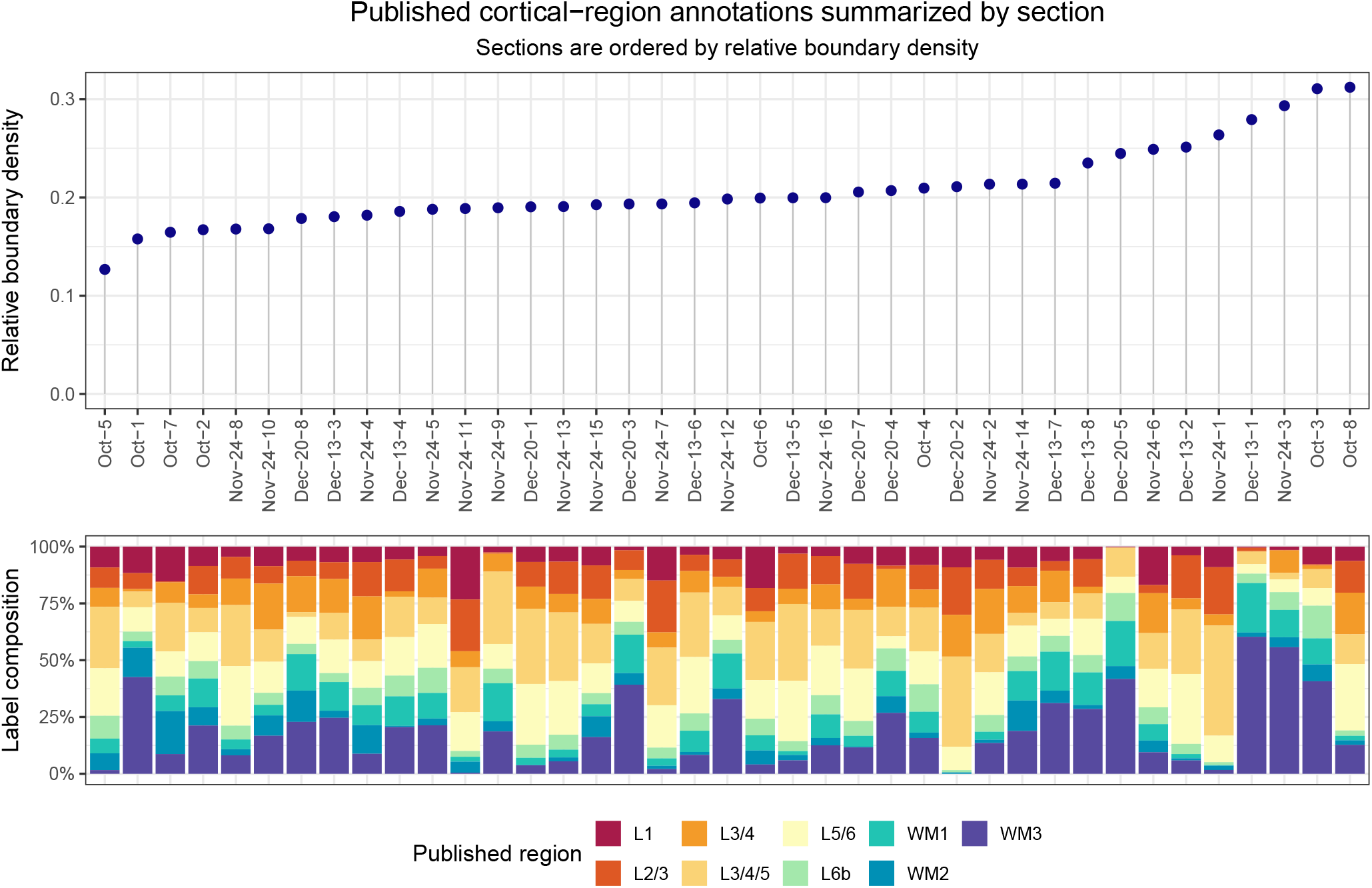
Relative BD and label composition across Miyoshi frontal-cortex sections. Top: sample-level relative BD for the published nine-region BayesSpace annotation across 39 sections, ordered by increasing value. Bottom: fractions of spots assigned to each region in the same section order. Section identifiers are abbreviated by omitting 2021 and Human/VisiumHuman and replacing underscores with hyphens. BD was calculated using *k* = 6 on the transformed Visium lattice.

**Supplementary Figure S13:**
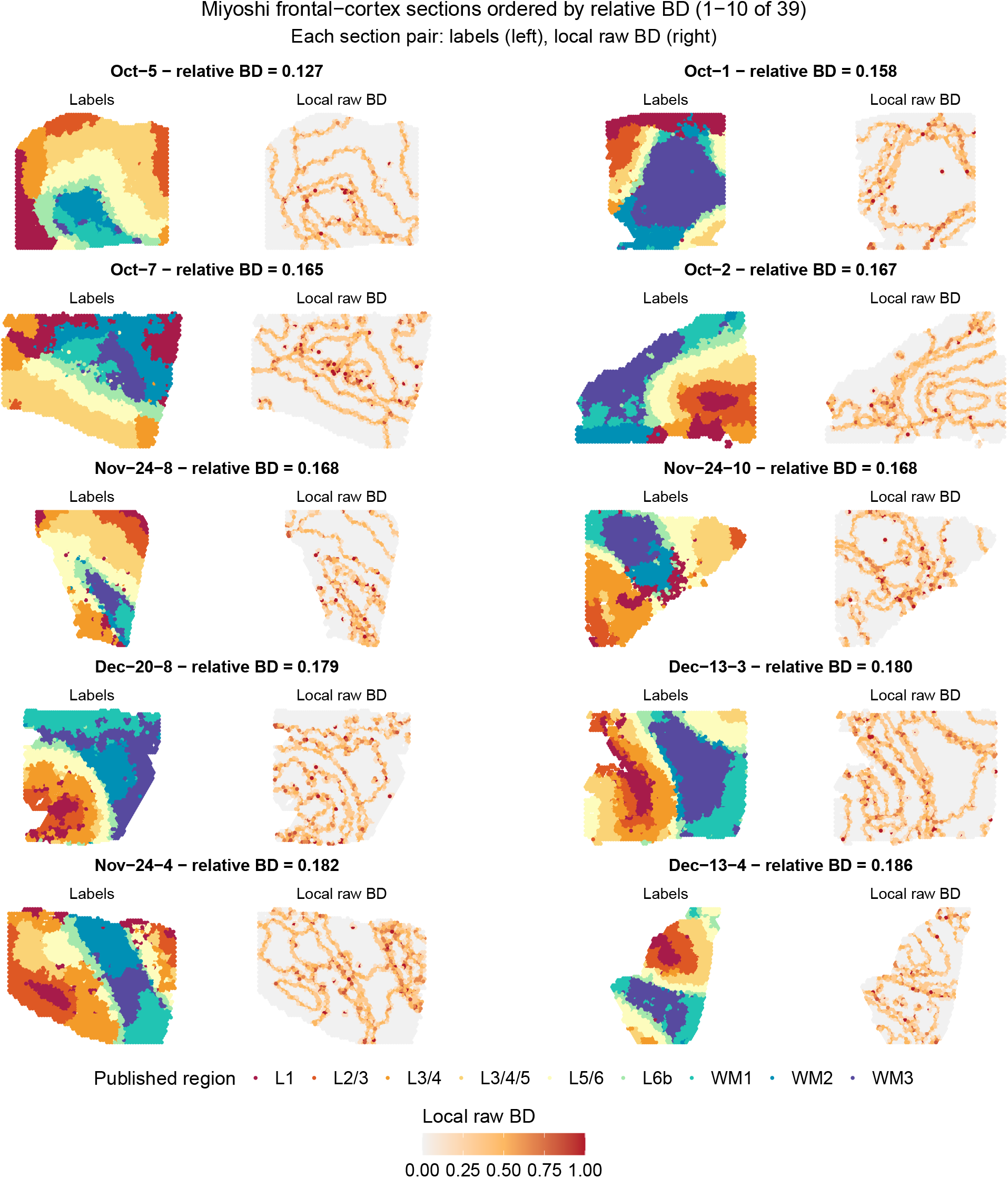
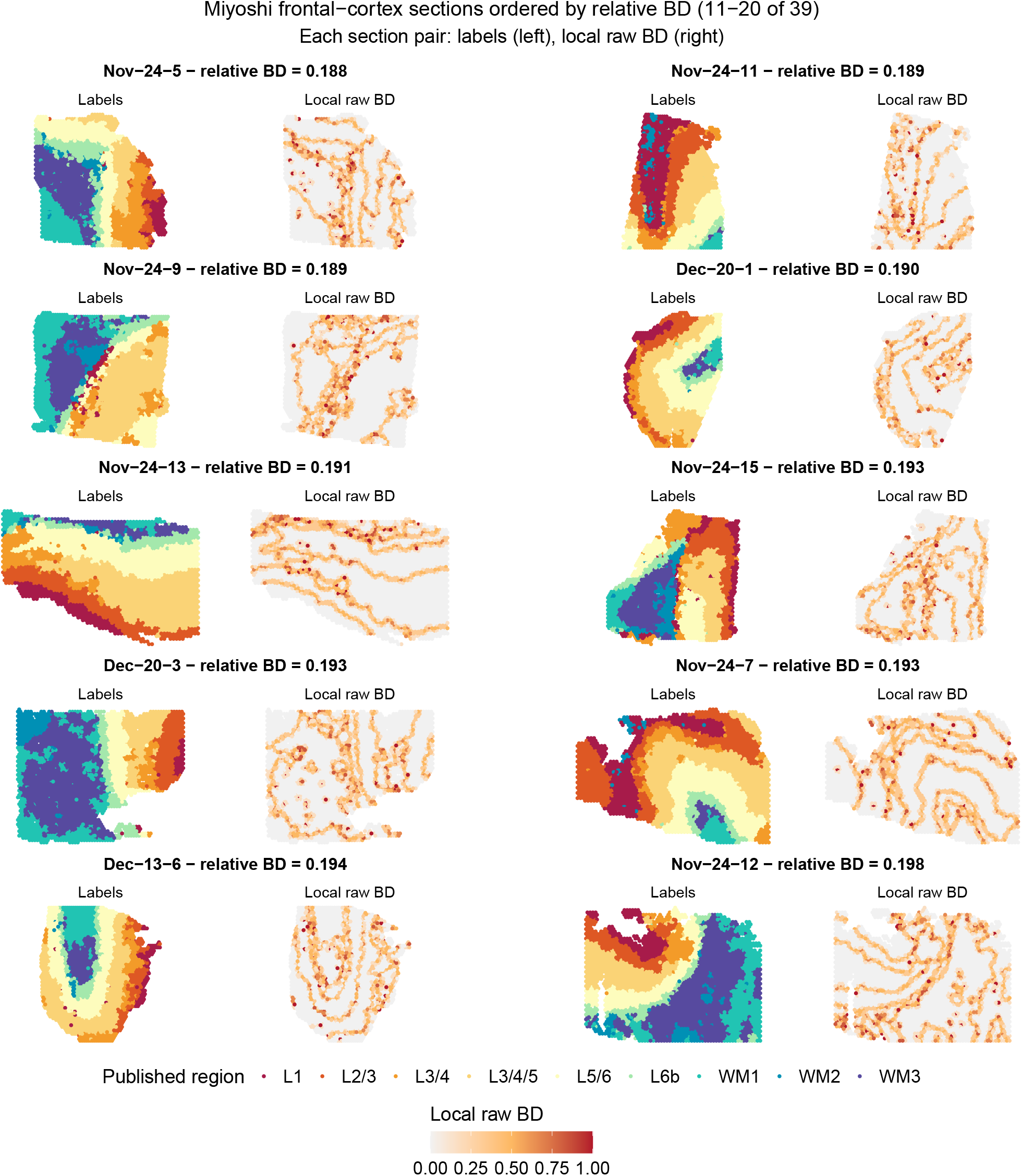
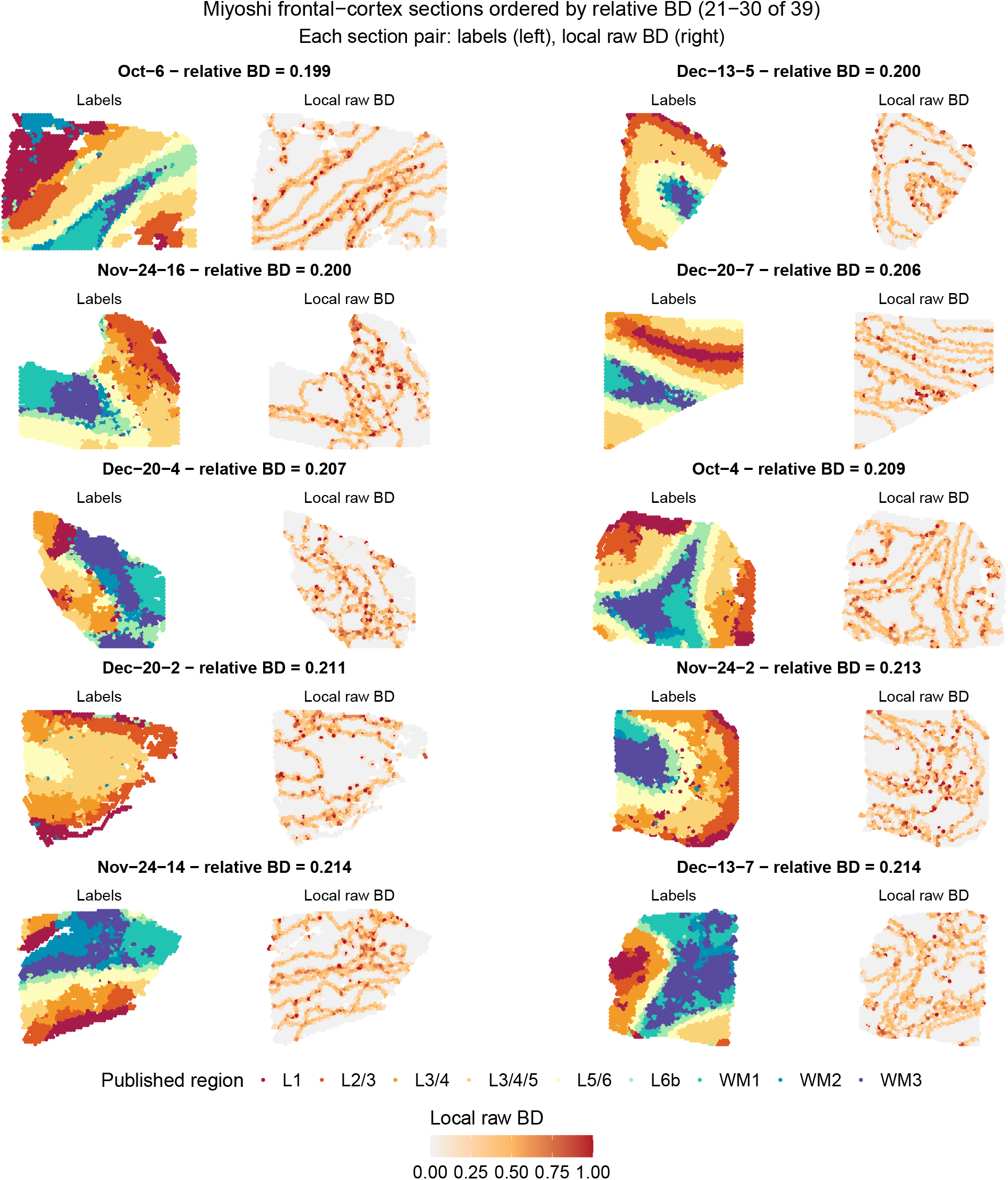
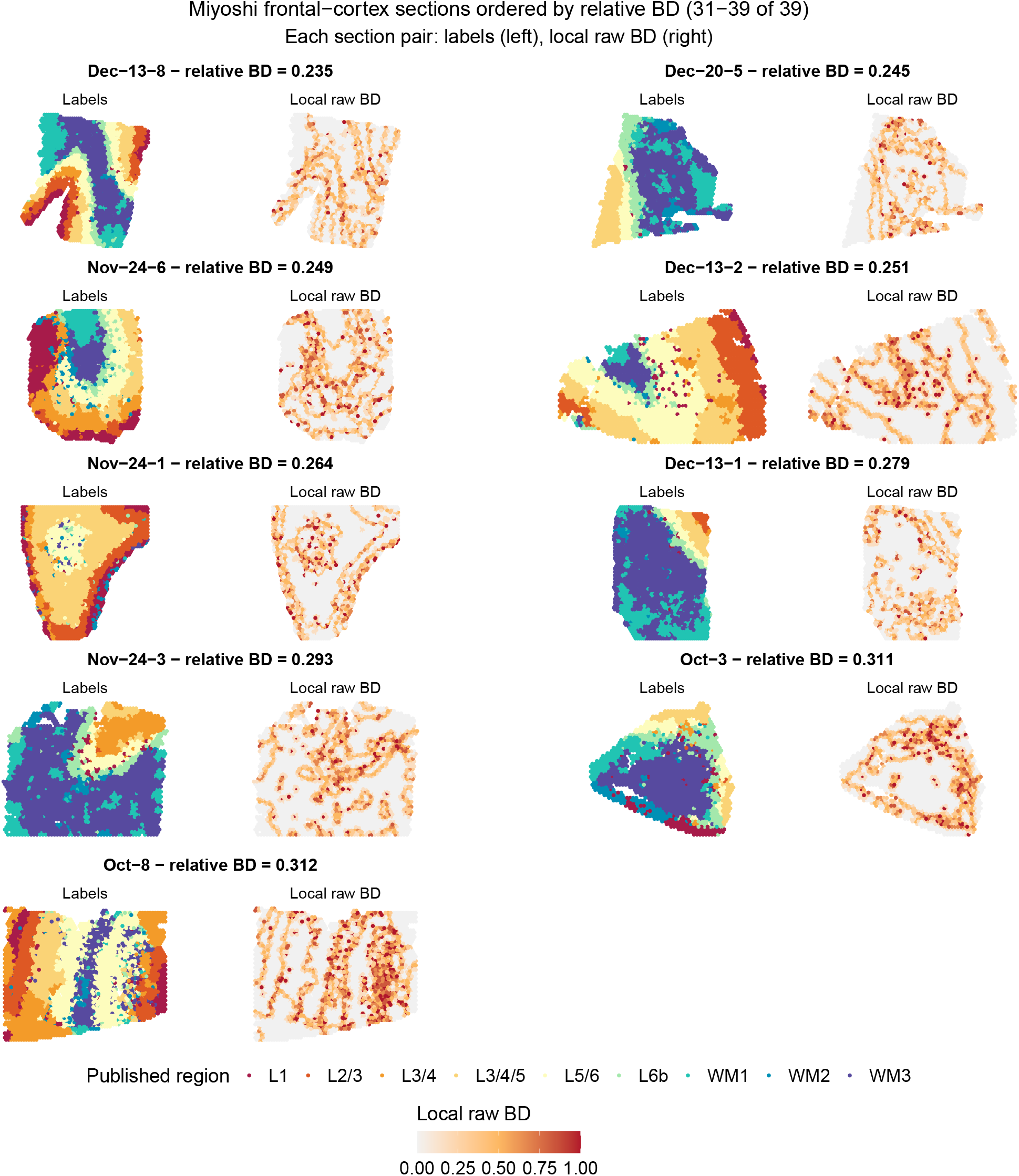
Published cortical-region annotations and local raw BD across all 39 Miyoshi frontal-cortex sections. Sections 1–10 of 39, ordered by sample-level relative BD for the published nine-region BayesSpace annotation. Read each row from left to right, with up to two section-pairs per row separated by a gutter. Each shared heading gives an abbreviated section identifier (omitting 2021 and Human/VisiumHuman) and sample-level relative BD. For each section, the left map shows the *q* = 9 annotation and the right map shows corresponding local raw BD calculated using *k* = 6. Each page has one annotation legend and one common 0–1 local raw-BD legend. These spatially informed labels provide an annotation-summary example.

**Supplementary Figure S14:**
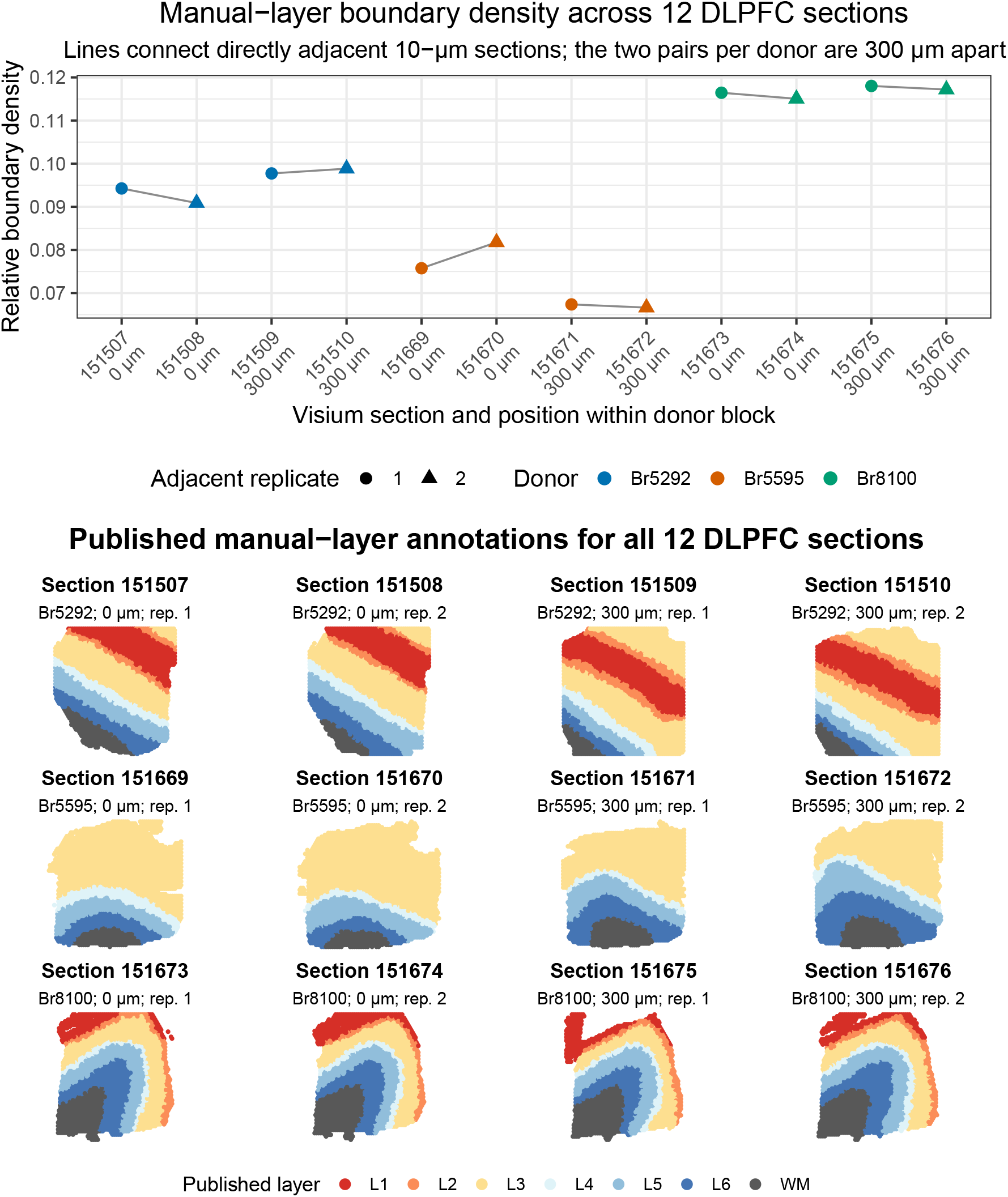
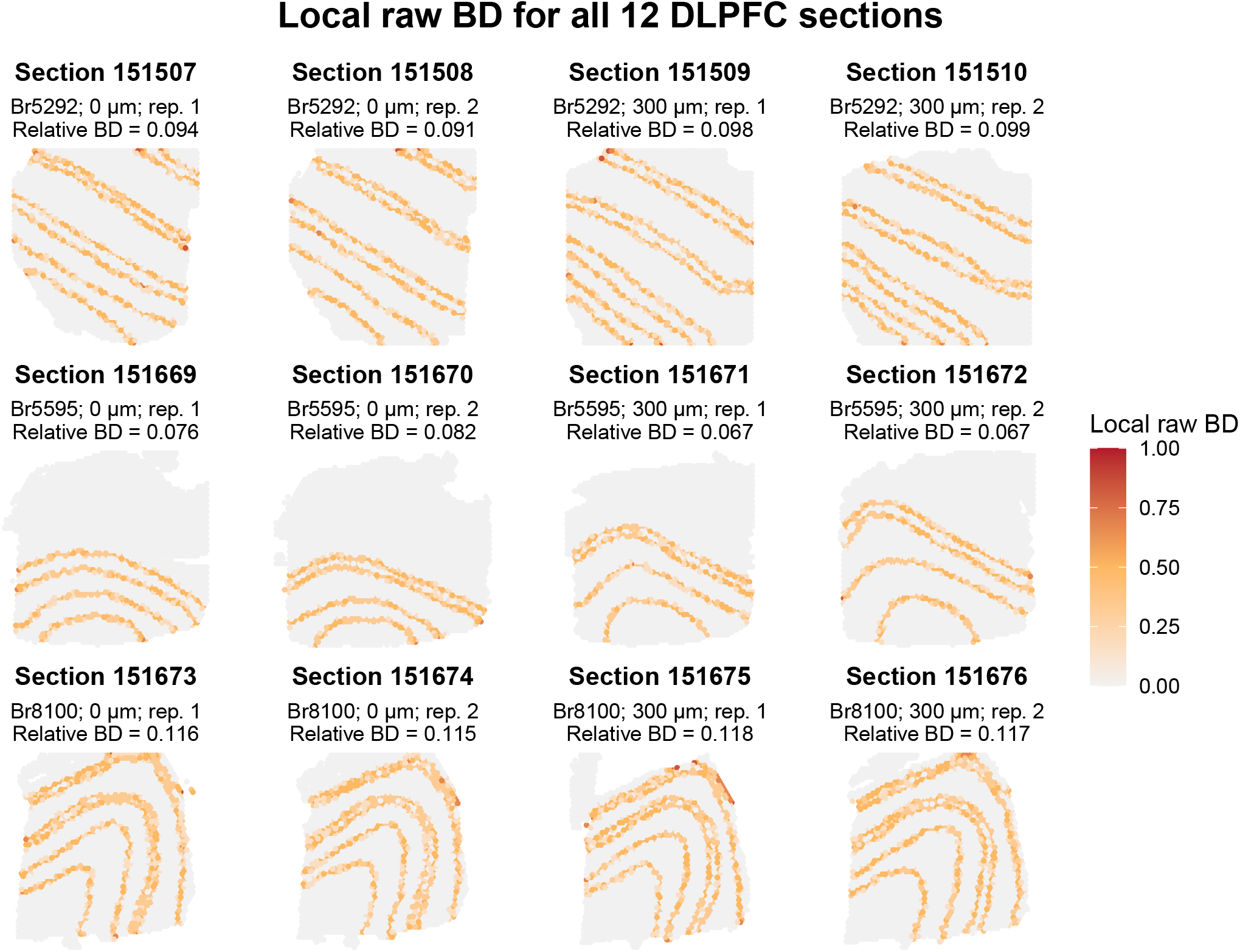
Relative BD and published manual annotations across 12 Maynard DLPFC sections. Top: relative BD for sections from three donors. Color denotes donor, shape distinguishes replicate 1 from replicate 2, and lines connect directly adjacent 10 *µ*m sections. Each donor contributed two adjacent pairs separated by 300 *µ*m. The comparison is descriptive and does not treat sections as independent replicates. Bottom: manual annotation maps. Rows correspond to donors Br5292, Br5595, and Br8100. Within each row, the first two columns form the adjacent pair at position 0 and the last two columns form the adjacent pair at 300 *µ*m in the tissue block. Colors denote layers L1–L6 and white matter. Br5595 lacked L1 and a distinct L2, and ambiguous L2/L3 spots were assigned to L3 in the original annotations; between-donor BD differences may therefore also reflect anatomical coverage and annotation granularity. The 352 spots without a manual layer assignment were excluded; all nonmissing manual labels were plotted, including 534 spots marked `discard=TRUE`in the deposited object.

**Supplementary Figure S15:**
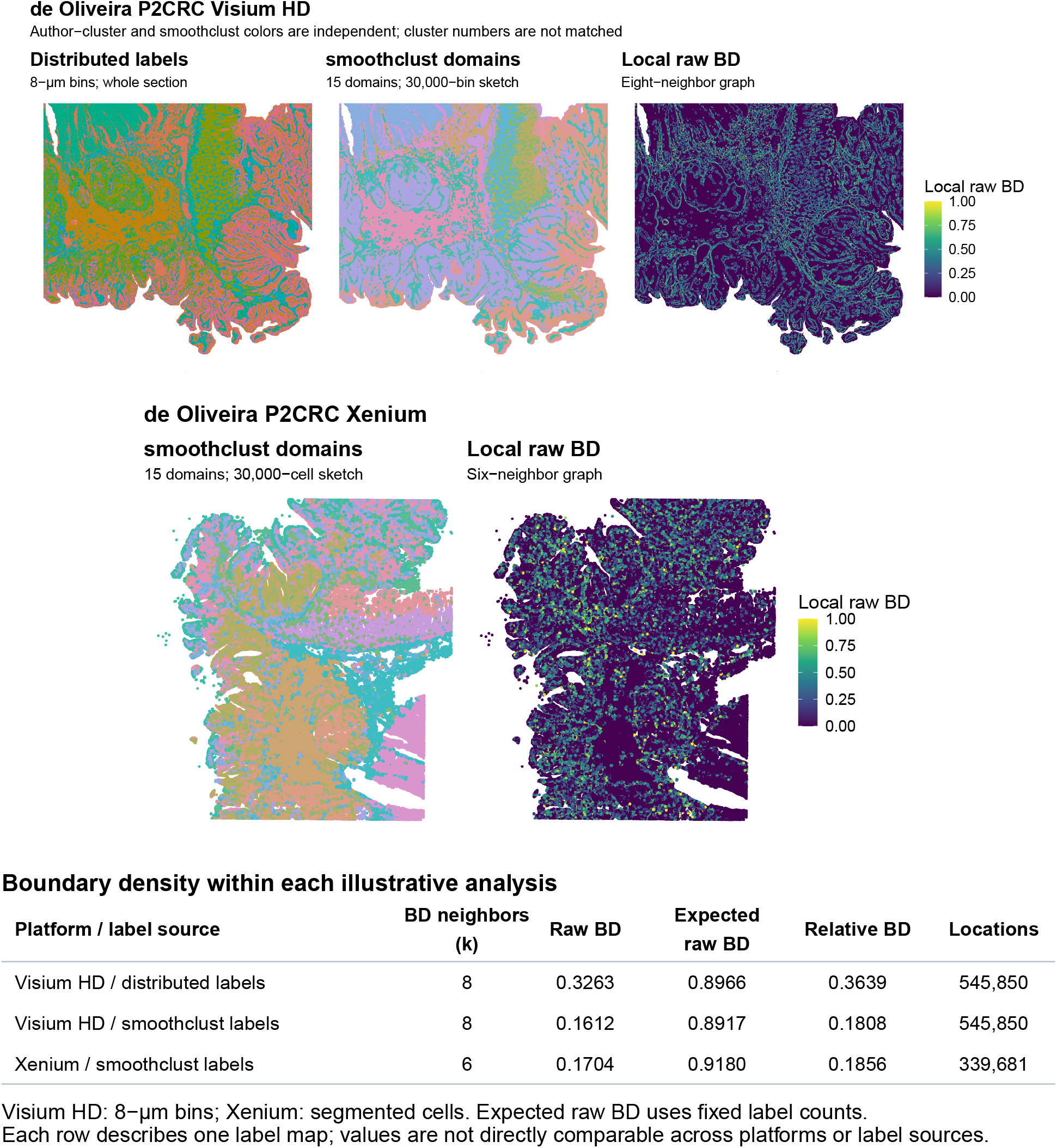
Illustrative full-section smoothclust and BD metric analyses on Visium HD and Xenium. Top: the complete P2CRC Visium HD section (545,850 nonempty 8 *µ*m bins), showing 15 cluster labels distributed with the source object, 15 smoothclust-derived domains, and local raw BD for these domains using *k* = 8 spatial neighbors. Colors and numeric identifiers are independent between label maps and do not imply one-to-one correspondence. Middle: a serial Xenium section from the same P2CRC FFPE tissue block (339,681 nonempty segmented cells), showing 15 smoothclust-derived domains and local raw BD using *k* = 6 cell neighbors. Both workflows used *k*_*s*_ = 18 smoothing neighbors on 1,000 selected genes for Visium HD and all 422 imported genes for Xenium, followed by a 30,000-location PCA/*k*-means fit and full-data projection and assignment. Bottom: neighborhood sizes, location counts, raw BD, expected raw BD, and relative BD for the three label maps. Expectations are analytic under random relabeling with fixed label counts; each row summarizes one map.

**Supplementary Figure S16:**
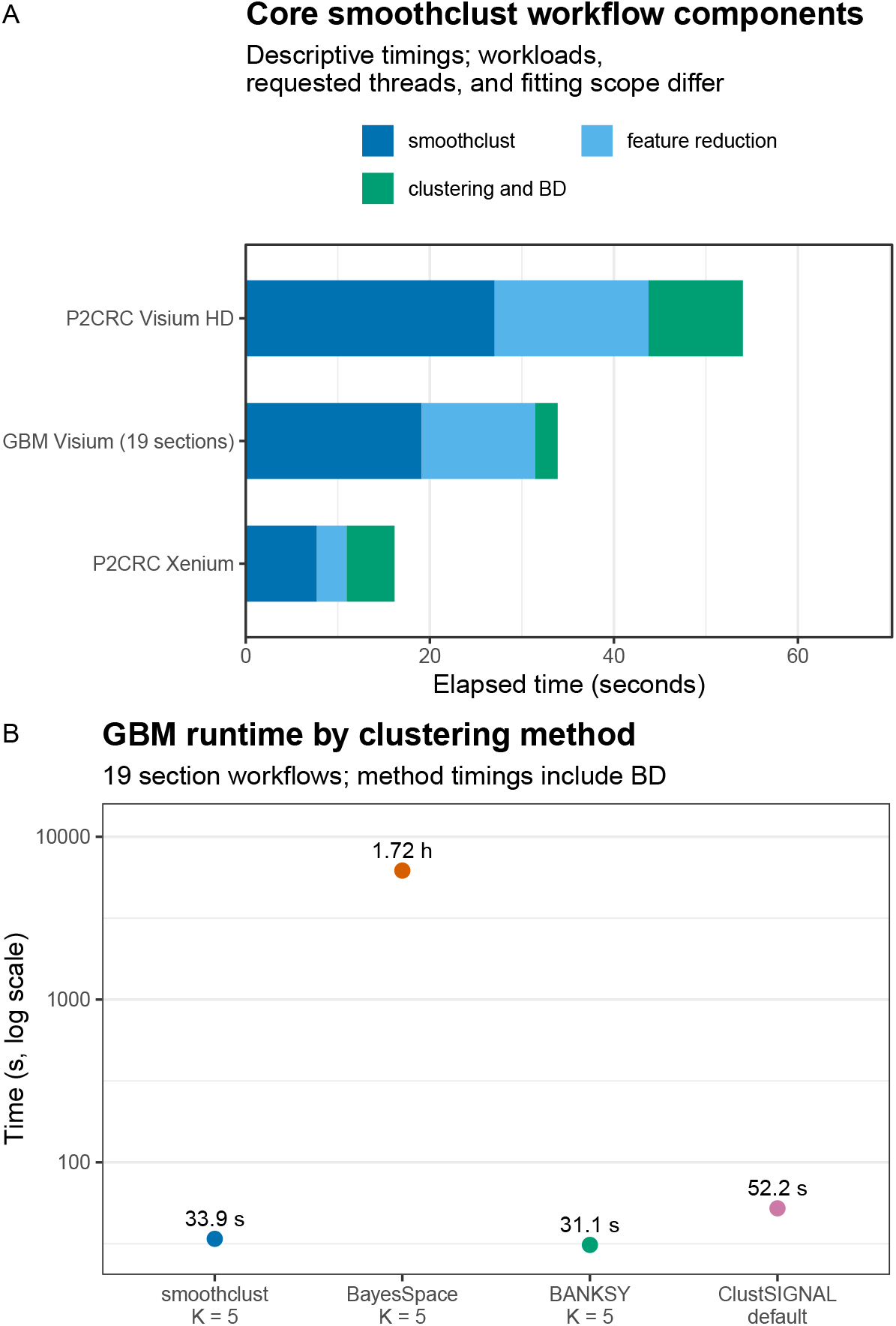
Core smoothclust workflow components and GBM method-comparison runtimes. (A) Stacked timings for smoothing, feature reduction, and clustering plus BD in the GBM Visium, P2CRC Visium HD, and P2CRC Xenium analyses, on a linear axis. The GBM total sums per-section *K* = 5 component timers; Visium HD and Xenium are directly observed single runs. Workflow sizes, fitting scope, timing exclusions, requested nearest-neighbor threads, and hardware are given in Supplementary Table S2. (B) Runtime comparison for smoothclust, BayesSpace, BANKSY, and ClustSIGNAL on the same processed GBM/glioma data, on a logarithmic axis. The first three used *K* = 5; ClustSIGNAL used defaults. The smoothclust time reuses the component-derived GBM total; external-method totals sum per-section timers. Method-comparison runtimes include method-specific preprocessing and dimension reduction where applicable, clustering or labeling, and the BD metric calculation; they exclude download, import, shared filtering, output, and plotting. GBM requested 14 threads where supported.

## Notes

### Competing Interest Statement

The authors have declared no competing interest.

